# Active DNA demethylation is upstream of rod-photoreceptor fate determination and required for retinal development

**DOI:** 10.1101/2025.02.03.636318

**Authors:** Ismael Hernández-Núñez, Alaina Urman, Xiaodong Zhang, William Jacobs, Christy Hoffman, Sohini Rebba, Ellen G Harding, Qiang Li, Fengbiao Mao, Andi K Cani, Shiming Chen, Meelad M Dawlaty, Rajesh C Rao, Philip A Ruzycki, John R Edwards, Brian S Clark

## Abstract

Retinal cell fate specification from multipotent retinal progenitors is governed by dynamic changes in chromatin structure and gene expression. Methylation at cytosines in DNA (5mC) is actively regulated for proper control of gene expression and chromatin architecture. Numerous genes display active DNA demethylation across retinal development; a process that requires oxidation of 5mC to 5-hydroxymethylcytosine (5hmC) and is controlled by the ten-eleven translocation methylcytosine dioxygenase (TET) enzymes. Using an allelic series of conditional TET enzyme mutants, we determine that DNA demethylation is required upstream of NRL and NR2E3 expression for the establishment of rod-photoreceptor fate. Using histological, behavioral, transcriptomic, and base-pair resolution DNA methylation analyses, we establish that inhibition of active DNA demethylation results in global changes in gene expression and methylation patterns that prevent photoreceptor precursors from adopting a rod-photoreceptor fate, instead producing a retina in which all photoreceptors specify as cones. Our results establish the TET enzymes and DNA demethylation as critical regulators of retinal development and cell fate specification, elucidating a novel mechanism required for the specification of rod-photoreceptors.

## INTRODUCTION

The generation of the diverse array of retinal cell types is orchestrated through the active regulation of cell fate specification from a single pool of multipotent retinal progenitor cells (RPCs)^1–3^. Retinal cell fate specification across development is modulated by dynamic changes in chromatin structure and gene expression patterns to facilitate the temporal specification of retinal cell fates. The past few decades of work have identified transcription factors and gene regulatory networks that bias the temporal specification of retinal cell fates. This includes both temporal competence factors (ATOH7, POU2F1, IKZF1, FOXN4, CASZ1 and the NFI transcription factors ^4–15^ and transcription factors that promote specification of individual cell fates (i.e. NRL)^16–18^. However, the exact mechanisms by which these transcription factors are expressed and their role in controlling developmental competence in RPCs remains unclear. We still do not comprehensively understand the specific mechanism by which a specified neuronal precursor commits to a terminal fate, including differentiation of major cell type classes (retinal ganglion cells, horizontal cells, amacrine cells, rod and cone-photoreceptors, bipolar cells, and Müller glia) and specification of the >120 mouse retinal cell subtypes ^19–23^.

Recent work has focused on how temporal and cell type-specific epigenetic modifications, including histone modifications, chromatin accessibility, and DNA methylation patterns bias chromatin remodeling and gene transcription for the specification of retinal cell fates ^24–27^. In particular, DNA methylation profiling has revealed temporally dynamic and cell-type specific DNA methylation patterns ^27–31^.

DNA methylation is established through the addition of a methyl group to the 5 position of cytosine residues [5-methylcytosine (5mC)] ^32–35^ and primarily functions to repress gene expression ^36–38^. Both enzymatic and passive processes regulate the methylation status of DNA. DNA methylation is promoted by de-novo methyltransferases (DNMTs), including DNMT1, DNMT3a and DNMT3b. Hypermethylation of promoters or enhancer sequences is correlated with reduced gene expression ^35,36^ and has an important role during development, aging, and disease ^33^.

However, many regulatory elements, including enhancers and promoters, undergo a transition from hypermethylation early in development to hypomethylation in mature cell types ^37,39–46^. DNA demethylation is regulated via three distinct mechanisms; 1) a passive, DNA replication-dependent manner whereby methylation is reduced by 50% after each round of DNA synthesis; 2) an active process mediated by the Ten-eleven translocation (TET) methylcytosine dioxygenases, TET1, TET2 and TET3 ^47–49;^ Figure 1A); or 3) conversion of 5mC to 5hmC by the TET enzymes and subsequent passive loss of 5hmC during DNA replication and cell division. The TET enzymes oxidize 5mC to generate 5-hydroxymethylcytosine (5hmC). 5hmC can be subsequently further oxidized by the Tet enzymes to produce 5-formilcytosine (5-fC) and carboxylcytosine (5-caC)^49^ which are then converted back to cytosine by Thymine DNA-Glycosylase (TDG) and the base-excision repair (BER) pathway (Figure 1A)^44,50–54^. In addition to being an intermediate during demethylation, 5hmC is also maintained as a stable mark that functions to control gene expression in a context-specific fashion through regulation of gene transcription, RNA splicing, or local control of histone modifications and chromatin remodeling ^55^. Recent evidence has highlighted an important regulatory role for 5hmC during both nervous system development and aging ^56,57^. However, to date, the nucleotide-specific localization and significance of 5hmC deposition in retinal cell fate specification remains undetermined.

**Figure 1.**
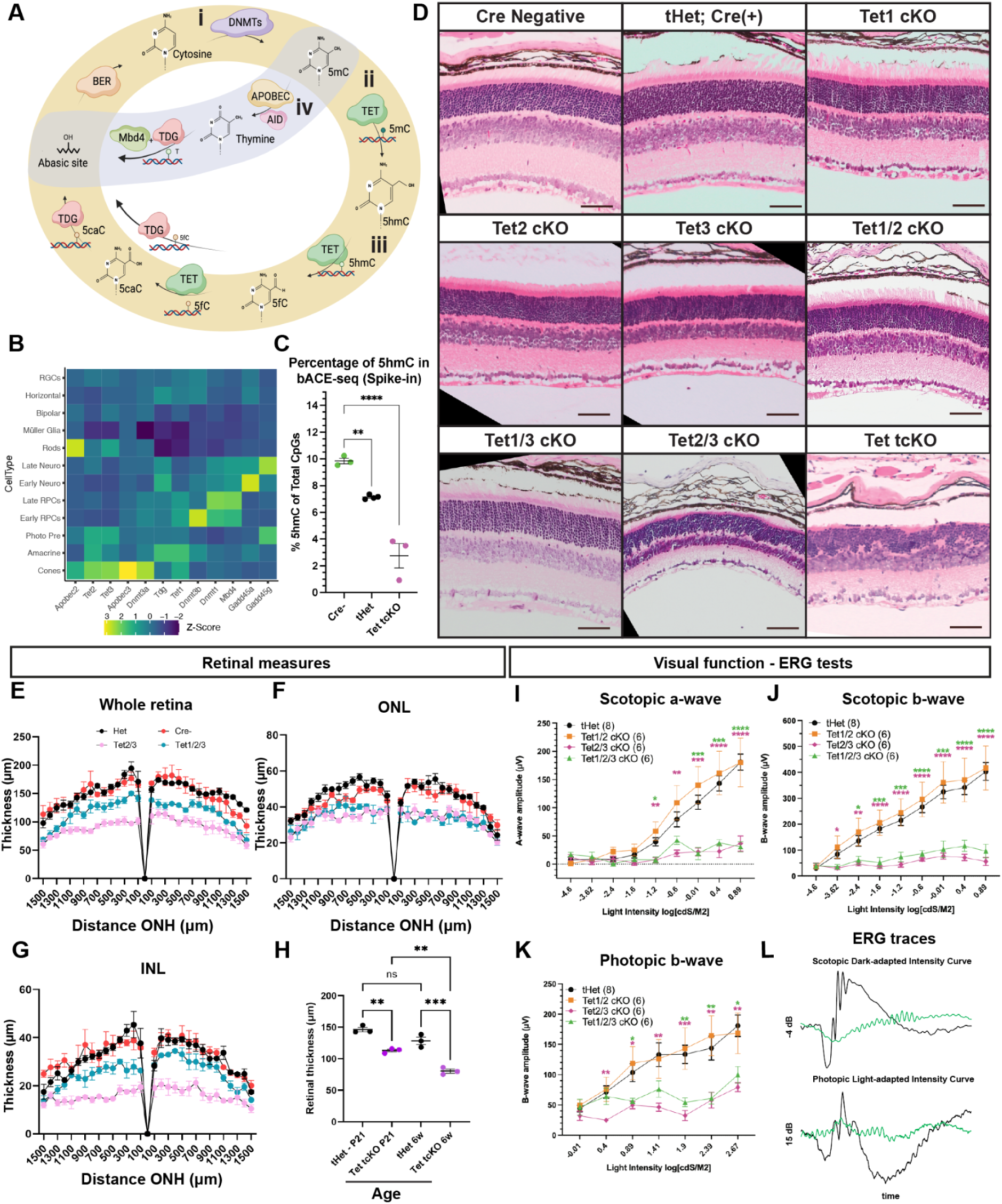
The Tet enzymes are required for retinal development and visual function. (A) Active DNA methylation cycle - i) 5mC is added by DNMTs. ii) The TET enzymes oxidize 5mC to 5hmC. iii) 5hmC is converted to 5fC and 5caC by the TET enzymes, followed by conversion back to cytosine by TDG and the base-excision repair pathway. iv) Alternatively, APOBEC converts 5mC to thymine, causing DNA mismatch. (B) Expression of DNA demethylation pathway components is enriched in photoreceptors. single-cell RNA-sequencing (scRNAseq) data from Clark et al. (2019). (C) bACE-seq quantification of 5hmC across P21 retinas in Cre-, tHet, and Tet tcKO animal models. Statistics represent results of a One-way ANOVA followed by a Dunnett’s Multiple Comparisons test (** - p < 0.01; **** - p < 0.0001). (D) H&E staining of an allelic series of TET conditional P21 mutants. (E) Whole retina, (F) outer nuclear layer (ONL) and (G) inner nuclear layer (INL) thickness measures at different eccentricities from the optic nerve head (ONH). Results display the mean + SEM for n=3 for each genotype in P21 retina. (H) Graph showing the comparison in the mean retinal thickness of control and Tet tcKO retinas between P21 and 6 weeks old retinas. Results display the mean + SEM for n=3 for each genotype. Statistics are the result of a Two-way ANOVA with, followed by a Tukey’s multiple comparisons test. ns: non significant; ** - p<0.01, *** - p<0.001; **** - p<0.0001. (I-K) Visual function testing as measured by full-field electroretinogram (ERG) indicating scotopic A-wave, scotopic B-wave and photopic B-wave amplitudes across different light intensities. Statistics are the result of a Two-way ANOVA with Geisser-Greenhouse correction, followed by Dunnett’s multiple comparison test to tHet controls. * - p<0.05; ** - p<0.01; *** - p<0.001; **** - p<0.0001. (L) Examples of ERG scotopic dark-adapted intensity (-4 dB) and photopic light-adapted intensity (15 dB) traces comparing tHet and Tet1/2/3 cKO retinas. Abbreviations: Neuro: Neurogenic RPCs; Photo Pre: Photoreceptor Precursor Cells; BER: base-excision repair; DNMTs: de-novo methyltransferases; TDG: thymine DNA glycosylase; Tets: tet-eleven translocation methylcytosine dioxygenases.

Genome-wide profiling of DNA methylation during retinal development has determined the temporal dynamics ^27^ and cell type-specific signatures of methylated DNA sequences ^28–31^, with 3-38% of genes displaying an inverse correlation of local DNA methylation and RNA transcript expression ^27^. Many cell type-specific gene loci undergo DNA demethylation as RPCs divide to generate post-mitotic cell types. Numerous gene promoters and gene bodies of rod and cone-photoreceptor genes are methylated in RPCs but display low DNA methylation levels and increased chromatin accessibility in mature photoreceptors ^27,29,30,58^. This has led to the hypothesis that active DNA demethylation plays an essential role in retinal cell fate specification and development ^31,58^. Supporting this hypothesis, alterations to retinal development, including improper specification of the eye-field were observed in Tet3 deficient *Xenopus* ^59^ or improper photoreceptor development in both Dnmt1/Dnmt3a/Dnmt3b conditional mutant mice ^60^ and Tet2/Tet3 mutant zebrafish ^61^ have been observed. Mouse models removing all TET enzymes exhibit early lethality during gastrulation ^62^, precluding studies examining retinal development. Therefore, to fully understand the significance of DNA demethylation and 5hmC on retinal cell fate specification and development in the absence of early eye-field patterning or systemic phenotypes, conditional mouse models for Tet1, Tet2, and Tet3 are required.

In this study, we utilize an allelic series of Tet enzyme conditional mouse mutants, removing the Tet enzymes within RPCs to determine the significance of active DNA demethylation and 5hmC for retinal cell fate specification and retinal development. Our studies indicate that most single and double Tet mutant combinations result in normal retinal morphology and visual function. However, Tet2/3 double and Tet tcKO retinas display abnormal retinal morphology, lack visual function, and display deficiencies in photoreceptor fate specification. Inhibition of the TET enzymes disrupts DNA demethylation and 5hmC production, preventing photoreceptors from differentiating into rod-photoreceptors and causing them to instead adopt a cone-like fate. Comprehensive transcriptional and single-base resolution 5mC and 5hmC analyses indicate substantial prevalence of 5hmC marks across the genome that are lost when the TET enzymes are removed. These data indicate that 5hmC likely plays a critical role in regulating the temporal DNA demethylation and activation of critical gene regulatory networks (GRNs) that regulate rod fate choice. Finally, we observe that TET3 expression and 5hmC modifications are both reduced in human retinoblastoma cells and compare the similarities in gene expression changes resulting from 5hmC loss in retinoblastoma to that of Tet tcKO mouse retinas. In combination, our work establishes active DNA demethylation as an initial regulator of rod fate choice, required for NRL and NR2E3 expression and the establishment of photoreceptor gene regulatory networks.

## MATERIAL AND METHODS

### Mice

Postnatal day (P) P0, P21 and 6 weeks old Cre-negative, heterozygous and Tet conditional knockouts (cKO) were generated using the Tg(Chx10-EGFP/cre,-ALPP)2Clc/J (*Chx10*::Cre-GFP, ^63^; Tet1*^loxp/loxp^*; Tet2*^loxp/loxp^*; Tet3*^loxp/loxp^* ^43^). All mice were housed in a climate-controlled pathogen-free facility on a 14/10 hour light/dark cycle. All experimental procedures were preapproved by the Washington University in St. Louis School of Medicine.

### Human Retinoblastoma samples

Approval of human studies was obtained by the University of Michigan Institutional Review Board under approval number PRO00011471 - Epigenetic regulation of ocular/orbital development, homeostasis and disease. Human retinoblastoma or unaffected control retinal samples were obtained through a retrospective search of the electronic pathology archives for cases of enucleation for retinoblastoma (RB) without prior treatment, diagnosed between January 1988 and June 2014. FFPE tissue blocks were cut into 5μm sections and stained with hematoxylin and eosin (H&E). Slides were reviewed and histologic features recorded in conjunction with review of the gross anatomical description by the ophthalmic pathologist as previously described ^64^. Portions of the slides with RB tumor and those containing normal retina without the presence of tumor involvement were used for downstream immunohistochemical and sequencing analyses.

### Tissue processing for H&E staining

Eyes were enucleated from euthanized animals and placed in 4% paraformaldehyde (PFA) for 1-2 minutes before generating a corneal tag by removing a portion of the ventral cornea to enable dorsal-ventral orientation following sectioning. Eyes are then fixed overnight in 4% PFA overnight followed by washing in phosphate buffered saline (PBS) for 3 minutes. Eyes are then placed in 70% ethanol at 4°C until paraffin embedding.

Embedding and sectioning of paraffin blocks and H&E staining was performed following standard protocols by the histology core in the Department of Ophthalmology and Visual Sciences at Washington University.

### Electroretinograms (ERGS)

ERGs were performed as previously described ^65^. Briefly, tests were performed on a visual electrodiagnostic system (UTASE3000 with EM for Windows; LKC Technologies, Inc., Gaithersburg, MD, USA) while mouse body temperature was maintained at 37°C ± 0.5°C with a heating pad controlled by a rectal temperature probe (FHC, Inc., Bowdoin, ME, USA). Pupils were dilated with 1.0% atropine sulfate (Bausch & Lomb, Tampa, FL, USA), and dilation and corneal hydration were maintained during testing by positioning the platinum wire loop recording electrodes in a mixture of atropine and 1.25% hydroxypropyl methylcellulose (GONAK; Akorn, Inc., Buffalo Grove, IL, USA). Mice were tested without knowledge of genotype. Bilateral flash ERG responses were obtained; the set of recordings displaying larger peak amplitudes were correlated with genotype information for statistical analyses. Differences in peak amplitude response at all light intensities were determined using a Two-way ANOVA and with Geisser-Greenhouse correction, followed by a Dunnett’s multiple comparison tests were performed using GraphPad Prism v10.2.3 (GraphPad Software, Inc., La Jolla, CA, USA).

### Immunohistochemistry

Eyes were enucleated from animals and placed in cold 4% paraformaldehyde for 1 hour followed by washing in 1X PBS. Retinas were then dissected to remove the choroid/sclera, RPE, and anterior segment and placed into 30% sucrose in PBS overnight at 4°C. Retinas were placed into 30% sucrose in PBS:OCT (1:1) overnight at 4°C and subsequently mounted in Tissue-Tek OCT media (VWR) for sectioning. Immunohistochemistry was performed following standard protocols. Briefly, slides are air dried and then washed in 1X PBS and placed into blocking solution [(1X PBS, 5% horse serum, 0.2% triton, 0.02% sodium azide, 0.1% bovine serum albumin (BSA)] for 2 hours. Slides are placed in the primary antibody diluted in blocking solution overnight at 4°C (Table S1). Slides are then washed in 1X PBS plus 0.05% triton three times for 5 minutes each. Primary antibodies are detected through incubation using fluorescently-tagged secondary antibodies diluted 1:500 in blocking buffer for 2 hours in the dark. Slides are then washed in 1X PBS plus 0.05% triton and then nuclei are counterstained using DAPI (1:3000 in 1X PBS plus 0.05% triton). Slides are then coverslipped using Vectashield HardSet Antifade Mounting Medium (Vector Labs).

### EdU experiments

Newborn (P0) mice were injected with EdU (10µM final concentration) and retinas were harvested at P1 or P14. Retinas are then processed for immunohistochemistry. EdU staining was performed using the Click-IT EdU Alexa Fluor 647 imaging kit (Invitrogen) following manufacturer’s instructions, with slides placed into blocking steps for the immunohistochemistry protocol directly after EdU detection. Nuclei were counterstained with DAPI (1:3000) and coverslipped using Vectashield (Vector Labs).

### Cell counts and statistics

For measuring the retinal thickness, measurements were taken from 1500µm both dorsally and ventrally from the optic nerve head (ONH). For cell counts, three images per retina were used to generate an average for each individual replicate. Replicate samples are from independent animals. Cell counts of control and Tet mutant retinas were made from a 200 x 200 µm area. Cell proportions were calculated by determining the number of marker positive cells divided by the total DAPI+ nuclei within the region of interest. Cell proportions in EdU experiments were calculated by determining the number of double positive cells (marker+/EdU+) divided by the total number of EdU+ cells in the whole image. Measures of retinal layers thickness, photoreceptor layers, and cell counts were performed using Fiji software 1.54f ^66^. Statistical analyses were performed with GraphPad Prism v10.2.3 (GraphPad Software, Inc., La Jolla, CA, USA).

### Imaging

Immunohistological data from H&E staining was imaged and photographed on the Zeiss Axio Observer inverted microscope coupled with an Axiocam 208 color camera (Zeiss). Immunohistochemical data from fluorescence labeling was imaged and photographed using a LSM800 confocal (Zeiss). 63x microphotographs were taken using the Airyscan imaging tool. Figure preparation was performed using Adobe Illustrator (Adobe, San Jose, CA). Contrast and brightness were minimally adjusted using Fiji software 1.54f ^66^.

### Mouse Retina RNA and DNA isolation

RNA and DNA were isolated following a modified protocol from TRIZOL Reagent isolation protocol. Briefly, retinal samples were placed in 1mL TRIZOL solution and homogenized. 200µl of chloroform was added to homogenized TRIZOL solution, mixed by vortexing and incubated for 3 min at RT. Lysates were centrifuged for 10 min at 4°C at max speed (16,000g). The aqueous phase with RNA was transferred into a fresh tube without disturbing the interphase. The rest of the volume, containing DNA and proteins was washed with 300µl 100% ethanol, inverted several times, and then centrifuged for 5 min at 4°C at 2000g. The supernatant containing proteins was carefully removed so as to not disturb the precipitated DNA pellet.

### Mouse Retina RNA extraction

RNA was extracted following standard protocol of the RNA Clean & Concentrator-5 (Zymo Research). Briefly, an equal volume of 100% ethanol was added to the aqueous phase containing RNA from the previous step. Samples were transferred to Zymo-Spin IC Column in a collection tube and centrifuge at 16000g. After discarding the flow-through, an in-column DNAse I treatment was performed. 400µl of RNA Prep buffer was added to the column and centrifuged at 16000g. After discarding the flow-through, two subsequent washes with RNA wash buffer were made. Then, the column is transferred into an RNAse-free tube and RNA eluted from the column using 30µl DNase/RNase-free water. RNA concentrations were measured using a Qubit fluorometer.

### Mouse Retina DNA extraction

After discarding the supernatant containing proteins, the DNA pellet was washed in 1mL of 100mM sodium citrate in 10% ethanol and incubated for 30 min at RT followed by centrifugation 5 min at 4°C at 2000g and removal of the resulting supernatant. This step is repeated twice. 1.5mL of 75% ethanol was added to the DNA pellet and incubated for 20 min at RT using occasional mixing during the 20-minute incubation. The supernatant is discarded, and the DNA pellet is air dried for 10 min. 100µl of 8mM NaOH in 1mM EDTA pH 7-8 was added to resuspend the DNA pellet. After that, samples are centrifuged for 10 min at 4°C and max speed (16,000g). The supernatant is transferred to a new tube and stored at -20°C. DNA concentrations were measured in a Qubit fluorometer.

### Nuclei Isolation and single nucleus RNA-sequencing (snRNA-seq) library preparation

Dissected retina tissue was dissociated on ice in 500ul cold Homogenization Buffer (25mM KCl, 5mM MgCl2, 10mM Tris-HCl, 1mM DTT, complete Mini Protease Inhibitor Cocktail (Roche)) with RNase inhibitors (200U/mL RNasin (Promega), 200U/mL SUPERaseIn (Invitrogen)) using a Dounce homogenizer, with 15 strokes each of pestles A and B. The lysate was strained through a 40um cell strainer, which was then rinsed with 9mL additional Homogenization Buffer. The strained lysate was centrifuged for 5 min at 500xg in a 4°C swinging-bucket centrifuge. After discarding the supernatant, the nuclei pellet was gently resuspended in 1mL cold Resuspension Buffer (25mM KCl, 3mM MgCl2, 50mM Tris-HCl, 1mM DTT) with RNase inhibitors (200U/mL RNasin (Promega), 200U/mL SUPERaseIn (Invitrogen)) using a wide-bore pipet tip. Nuclear concentration was quantified by Countess. Appropriate numbers of nuclei were then processed according to the PIPseq T20 3’ Single Cell RNA Kit v4.0 (Fluent BioSciences). Completed libraries were sequenced on a NovaSeq X Plus (Illumina Inc) and reads processed using pipseeker (v3.3.0) and aligned to the GRCm39 reference genome. Libraries were sequenced on the Illumina NovaSeq X Plus using 2 x 150bp paired-end reads.

### bACE-seq library preparation

bACE-Seq libraries were made using the protocol from ^67^ with minor modifications. In brief, 10-15ng of RNAse treated, TRIZOL purified gDNA was used for each sample. Samples were diluted to 10µL in MilliQ water and pre-heated by incubating at 50C for 20 minutes on a thermocycler with a heated lid set to 95°C. 32.5µL of CT Conversion Reagent (Zymo EZ DNA Methylation Direct Kit) was then added to each sample before running the the following program on a thermocycler (98°C for 8min, 64°C for 105 min, followed by 98°C for 8 min, 64°C for 105 min), with the lid set to 95°C. Following the bisulfite reaction, the tubes were placed at -20°C overnight, or for 16 hours. The bisulfite converted DNA was then purified using the Zymo EZ Methylation Direct Kit, eluting with 17µL of MilliQ water for a final elution volume of 16µL.

Each sample was incubated at 95°C for 3 minutes on a thermocycler, with the lid set to 105°C. The samples were then immediately placed into a dry-ice ethanol bath to snap cool them and were left in the cooling bath for 5 minutes. The samples were then placed on a pre-chilled PCR rack and kept on ice. 8µL of premixed APOBEC reaction mixture (2.4µL APOBEC reaction buffer (E7134A, NEB), 0.48µL APOBEC (E7133AA, NEB), 0.48µL BSA (B9000S, NEB), water to 8µL) was added to each tube and samples were incubated at 37°C on a thermocycler with the lid set to 95°C. After 30 min, the samples were mixed and placed back on the thermocycler for two and a half hours at 37°C with the lid set to 95°C. DNA was purified from the reaction using 2X Agencourt AMPure XP SPRI beads (add 2X volume of SPRI beads, incubate at room temperature for 10 minutes, place samples on magnet and wash 2 times with fresh 80% ethanol, dry for 3 min, elute in 9µL Low EDTA TE (IDT).

Sequencing libraries were constructed using the xGen Adaptase Module with minor modification. 250nM of random stubby index primer was added to each sample (xGen Adaptase Module, P5L_AD002_H). Samples were incubated at 95°C on a thermocycler with the lid set to 105°C for 3 min, then cooled on ice for 2 min. 10µL of BST-based random priming mixture (2µL 10X ISO AMP Buffer II (B0374S, NEB), 0.8µL BST 3.0 (M0374S, NEB), 1.2µL 200mM MgCl2, 2.8µL 10mM dNTP, 3.2 µL water) was then added to each sample and incubated at 60°C for 1 hour on a thermocycler with the lid set to 95°C. Fragment ends were blunted and dephosphorylated by adding 2µl Exonuclease I (M0293L, NEB) and 1µl shrimp alkaline phosphatase (M0371S, NEB) to each sample. The samples were then incubated at 37°C for 30 min and purified using a 1.6X SPRI beads (add 1.6X volume of SPRI beads, incubate at room temperature for 10 minutes, place samples on magnet and wash 2 times with fresh 80% ethanol, dry for 3 min, elute) Samples were eluted in 10 µL Low EDTA TE (IDT). In brief, samples were denatured by incubating at 95°C on a thermocycler with the lid set to 105°C for 3 min, then cooled on ice for 2 min. Samples were indexed and amplified using the PCR protocol in the xGen Adaptase Module using 21 cycles. The indexed samples were purified using 1.6X SPRI beads (add 1.6X volume of SPRI beads, incubate at room temperature for 10 minutes, place samples on magnet and wash 2 times with fresh 80% ethanol, dry for 3 min, elute) and eluted in 12µL Low EDTA TE (IDT).

Libraries were initially sequenced by spike-in low pass sequencing 2x150 bp on an Illumina MiniSeq (∼100k reads per sample) to ensure that libraries were high quality (reasonable mapping and Apobec conversion efficiencies) before being sequenced using 2x150 paired end reads on an Illumina NovaSeq X Plus.

### WGBS library preparation

Genomic DNA was quantified using the Qubit fluorometer. 200ng of gDNA, including 0.2% Lambda DNA (N6-methyladenine-free; NEB) was fragmented in a final volume of 50uL using the Covaris LE220 targeting ∼350bp inserts. A 1.5x AMPure clean-up was utilized after fragmentation to concentrate the sample. The fragmented gDNA was bisulfite converted with the EZ-96 DNA Methylation-Gold Mag Prep Kit (Zymo Research, Cat # D5043) according to the manufacturer’s recommendations. Bisulfite converted DNA was quantitated on the Qubit Fluorometer using the ssDNA Assay Kit (Thermo Fisher Scientific, Cat #Q10212). Whole genome bisulfite libraries were constructed with ∼100ng of converted DNA using the xGen™ Methyl-Seq Library Prep Kit (Integrated DNA Technologies, Cat # 10009825) using unique dual indexes (IDT, 0008053) and eight PCR cycles for incorporation and amplification of indexed libraries, followed by a final 0.85x AMPure cleanup. Final libraries are quality checked for average library size and concentration. Molar concentration of libraries was determined using the KAPA Library Quantification kit (Roche Diagnostics). Libraries were sequenced on NovaSeq X using 150bp paired end reads.

### RNA-seq library preparation

#### Mouse

For RNA-seq library preparation, SMARTer Stranded Total RNA High Input (RiboGone Mammalian) (Takara) was used. Briefly, extracted RNA first undergoes ribosomal RNA removal. Then, we proceeded with cDNA synthesis and purification. Following, RNA-seq library was amplified by PCR. Total RNA was captured and purified from RNA samples using Illumina TruSeq stranded primers. RNA-libraries concentrations were measured in a Qubit fluorometer and using the Agilent 2100 Bioanalyzer. 9 libraries were pooled and sequenced using the NovaSeq X Plus 300 cycles system with ∼300 million paired-reads per run, resulting in between 40 million and 55 million reads per library.

### Human Retinoblastoma

For each specimen, 3–7 10μm formalin-fixed, paraffin-embedded sections were cut from a single representative block per case, using macrodissection with a scalpel as needed, to enrich tumor content. RNA was isolated using the Qiagen Allprep formalin-fixed, paraffin-embedded DNA/RNA kit (Qiagen, Valencia, CA) and quantified as previously described ^64^. Library preparation and sequencing was performed using TruSeq® RNA Access Library Preparation Kit (Illumina) and Illumina HiSeq 4000 Libraries platform at University of Michigan Sequencing Core.

### Sequencing Analysis

#### bACE-seq

Data were processed similarly to ^67^ with minor modification. Paired-end sequencing reads were processed with TrimGalore (v0.4.4_dev; https://github.com/FelixKrueger/TrimGalore) to remove adapters and low-quality sequences using the parameters --paired --clip_R1 16 --clip_R2 16. The trimmed reads were independently aligned to the mouse genome (mm10) using Bismark (v0.18.2). Read 1 (R1) was aligned with Bowtie2 in PBAT mode (--pbat flag) and the following settings: -q --score-min L,0,-0.2 -p 4 --reorder --ignore-quals --no-mixed --no-discordant –dovetail --maxins 500 --directional. Read 2 (R2) was mapped using Bowtie2 with default parameters. Aligned reads were filtered to retain only those with a mapping quality score ≥ 10 using Samtools (v1.15.1) and PCR duplicates were removed using Picard’s MarkDuplicates tool (GATK v4.1.3). CpG methylation levels were extracted with MethylDackel (v0.6.1; https://github.com/dpryan79/MethylDackel) using the --CHG and --CHH flags alongside default settings. Finally, strand-specific read counts were combined using a custom Python script.

#### WGBS

WGBS data were processed using the same pipeline as described for bACE-seq analysis (above), with modifications to the alignment steps. Reads were aligned to the mouse genome (mm10) using Bismark (v0.18.2)^68^ with default alignment parameters (PBAT mode turned off).

#### Differentially Methylated Region (DMR) Analysis

DMRs were identified using DSS (v2.48.0)^69^. The DMLtest function was applied with smoothing=TRUE and default parameters. DMRs were called with a p-value threshold of 0.01. To define DMRs, a minimum length of 50 bp and at least 3 CpG sites were required, with DMRs within 50 bp merged (following DSS default settings). DMR annotation was performed using ChIPseeker (v1.36.0)^70^, with the transcription start site (TSS) Region set to +/- 2000 bp and default settings. Mouse gene annotations from TxDb.Mmusculus.UCSC.mm10.knownGene were used for the annotation process.

### bACE-seq and WGBS-seq Data Integration

For each CpG site, 5mC and 5hmC levels were estimated using MLML, which calculates maximum likelihood estimates by integrating data from bACE-seq (5hmC) and WGBS-seq (5mC + 5hmC). MLML^71^ was applied with a significance threshold of α = 0.05 for the binomial test at each CpG site, and similarly for CpH sites in non-CpG analyses.

### RNAseq

Quality control of raw sequencing reads was performed using FastQC (v0.11.9) (https://www.bioinformatics.babraham.ac.uk/projects/fastqc/?utm_source), with summaries generated by MultiQC (v1.19)^72^. Adapter trimming and removal of low-quality bases were conducted using Trim Galore (v0.6.7) (https://www.bioinformatics.babraham.ac.uk/projects/trim_galore/). Trimmed reads were aligned to the GRCh38 or mm10 reference genomes, using STAR (v2.7.0f; with default parameters optimized for paired-end data, followed by post-alignment processing, including sorting and indexing of BAM files, using Samtools (v1.9)^73^. Gene-level quantification was performed using HTSeq (v0.11.2)^74^, and genome-wide read coverage was visualized using deepTools (v3.5 ^75^). Differential expression analysis was conducted in edgeR (v4.2.2)^76^ within R (v4.4.1), employing TMM normalization and statistical modeling, with significance thresholds set at |log2FC| ≥ 1 and FDR < 0.01. Heatmaps were generated using pheatmap (v1.0.12) https://CRAN.R-project.org/package=pheatmap), and Gene Ontology (GO) enrichment analysis was performed with clusterProfiler (v4.12.6)^77^.

#### Ortholog Mapping and Cross-Species Analysis

Human-mouse and mouse-human orthologs were identified by mapping Ensembl gene IDs between species using biomaRt (2.60.1)^78^. Differentially expressed genes were classified as upregulated, downregulated, or unchanged based on thresholds of |log2FC| ≥ 1 and FDR < 0.01, with orthologous genes analyzed using the same thresholds. Comparative boxplots were generated using ggplot2 (v3.5.1; https://ggplot2.tidyverse.org/)^79^, and Wilcoxon rank-sum tests were performed to evaluate gene expression differences across orthologs.

#### snRNA-seq analysis

To lessen the effects of ambient RNA contamination, resulting matrices from pipseeker outputs were processed using Cellbender v.0.3.0 ^80^. The priors used for the --expected-cells and --total-droplets-included parameters were 1000 and 350,000, respectively. A “full” model was used for both samples with the model dimensions set to --z-dim 256 and --z-layers 2048. Each model was trained to 150 epochs using a learning rate of 1x10^-7. Only cells with a 0.99 probability were used in the downstream analysis.

Resulting matrices from Cellbender were imported into Monocle3^81,82^ for single-cell analysis. Cells containing greater than 500 and less than 10000 transcripts with less than 20% mitochondrial transcripts were utilized for downstream analyses. Dimension reduction was performed using a preprocessed matrix with PCA followed by UMAP dimension reduction using the first 16 PCA dimensions. Cell type annotations were performed using marker gene expression within clusters for known marker genes for retinal cell types as determined previously ^14^. Differential expression analysis by genotype was performed using the Monocle3 fit_models function performed on all genes expressed within at least 100 cells and a q-value threshold of 1e-20.

## RESULTS

### Differential transcript enrichment of Tet transcripts within the developing retina

The cyclical process of DNA methylation/demethylation is regulated by numerous enzymes that facilitate cytosine modifications and DNA repair (Figure 1A). Our transcriptional profiling of single-cells across both mouse^14^ and human^83^ retinal development identified numerous components of the DNA methylation/demethylation pathways as differentially enriched across retinal development, including transcripts encoding the DNA methyltransferases (DNMT1, DNMT3A, DNMT3B) and the ten-eleven translocation methylcytosine dioxygenases (TET1, TET2, and TET3; Figure 1B). Prior work has indicated that the DNMTs are required for proper photoreceptor development ^84,85^; however the significance of 5hmC and TET-mediated DNA demethylation for retinal development and cell fate specification remains unclear ^31,61,86^. Tet1, Tet2 and Tet3 show differential transcript enrichment levels across retinal cell types, displaying enrichment within the photoreceptors, photoreceptor precursors, and bipolar cells (Figure 1B) ^61^.

### TET enzymes are required for retinal morphology, function, and cell fate specification

Recent work has highlighted the prevalence of 5hmC within the developing nervous system ^87,88^ and a requirement of DNA demethylation for glial fate specification ^89^. To determine if DNA demethylation and 5hmC deposition function similarly in the developing retina, we utilized conditional mouse knockout models targeting the TET enzymes, the key drivers of 5hmC. We chose to conditionally delete the TET enzymes (Tet1*^loxp/loxp^*; Tet2*^loxp/loxp^*; Tet3*^loxp/loxp^*) within developing mouse RPCs using the Tg(Chx10-EGFP/cre,-ALPP)2Clc/J (*Chx10*::Cre-GFP) transgenic line ^63^. To address mosaicism of the Chx10-Cre-GFP transgene ^63^, we implemented a breeding strategy involving Cre-positive incrosses to ensure two functional copies of the transgene. This approach mirrors strategies previously employed in Nfia/b/x triple mutant retina studies ^14^. Removal of the TET enzymes within RPCs allows passive, DNA replication dependent DNA demethylation to persist; however, TET-mediated active and passive DNA demethylation is inhibited in both proliferative and post-mitotic cells. We first validated the efficacy of our Tet loss-of-function strategy and inhibition of 5hmC using bisulfite-assisted APOBEC-coupled epigenetic sequencing (bACE-seq). Shallow sequencing of bACE-seq libraries in postnatal day 21 (P21) retinas from Cre-negative (Cre-), Tet1*^loxp^*^/+^; Tet2*^loxp^*^/+^; and Tet3*^loxp^*^/+^ triple heterozygous Chx10-Cre-(+) (tHet), and Tet1*^loxp^*^/loxp^; Tet2*^loxp^*^/loxp^; Tet3*^loxp^*^/loxp^ Chx10-Cre(+) triple mutants (Tet tcKO) indicated a Tet dosage-dependent decrease in 5hmC across genotypes (Figure 1C), highlighting effective knockout strategies.

The phenotypic effect of TET enzyme loss-of-function and reduced levels of 5hmC was first characterized on an allelic series of TET mutants using morphological analysis of H&E stained retinal cross-sections from 3-week old mice (postnatal day 21 - P21). We observed that the TET enzymes are individually dispensable for retinal development (Figure S1A). Morphological characterizations of Tet1/2, and Tet1/3 double mutants were also indistinguishable from Cre- and tHet controls (Figure 1D-E). However, RPC-specific deletion of Tet2/3 or all three TET enzymes (Tet tcKO) resulted in abnormal retinal development, characterized by an absence or significant attenuation of elongated photoreceptor outer segments, disruption of the outer plexiform layer (OPL) and a decrease in retinal thickness (Figures 1D-E; Figure S1A). This decrease in retinal thickness resulted from a decrease in the thickness of both the outer nuclear layer (ONL) and inner nuclear layer (INL; Figures 1D-H; Figure S1A). Quantification of the number of rows of photoreceptor nuclei suggests a loss of photoreceptor number or developmental disorganization in Tet tcKO retinas (One-way ANOVA followed by a Tukey’s comparisons test p = 0.0003; Brown-Forsythe test p = 0.8006; Figure S5B). Morphological characterizations of retinas at 6-weeks showed a continued thinning of retinas and persistent lack of photoreceptor outer segment morphogenesis (Figure 1H; Figures S1B-C).

To determine the consequence of TET enzyme loss-of-function on visual function, we performed electroretinograms (ERGs) on 6-week old mice. Consistent with the observed morphological disruptions (Figures S1B-C), we identified a significant attenuation of visual function in Tet2/3 cKO and Tet tcKO mice. In both Tet2/3 and Tet tcKO animals, we observed a reduction in the dark-adapted A-wave, dark-adapted B-wave and light-adapted B-wave compared to heterozygous control animals (Figures 1I-L), indicative of disrupted rod- and cone-photoreceptor responses to light stimuli and a failure to transduce light signals from photoreceptors to bipolar cells. The combined morphological and functional results indicate the significance of Tet2 and Tet3 enzymes - and potential redundant functions - in retinal development, as neither Tet1/2 or Tet1/3 double mutant retinas displayed gross morphological or visual function deficit phenotypes.

To assess the cause of reduced visual function in Tet2/3 cKO and Tet tcKO mutant retinas, we first characterized the specification of retinal cell fates using immunohistochemistry. We determined that the localization and distribution of cell type proportions across the allelic series of TET mutant retinas for retinal ganglion cells (RGCs), horizontal cells, amacrine cells, bipolar cells, and Müller glia using the cell type markers RBPMS, CALB1, PAX6, VSX2, and LHX2, respectively (Figure 2; Figure S2). In general, the localization of all cell types appeared normal, as cell type markers stratified to known nuclear layers. Assessments of RGC numbers across the allelic series of TET mutant retinas did not result in any significant differences across genotypes in comparison to Cre-negative controls (Figures 2A-B; Figure S2; One-way ANOVA followed by a Dunnett’s multiple comparisons test p = 0.0199; Brown-Forsythe test p = 0.7402). However, we did observe significant differences in the numbers of Calb1+ horizontal cells, Pax6+ amacrine cells, Vsx2+ bipolar cells, and Lhx2+ Müller glia across genotypes (Figure 2; Figure S2). There was a significant loss of horizontal cells marked by CALB1 in all retinas harboring Tet3 loss-of-function (Tet3, Tet1/3, Tet2/3, and Tet tcKO retinas; Figures 2C-D; Figure S2; One-way ANOVA followed by a Dunnett’s multiple comparisons test; p <0.0001; Brown-Forsythe test p = 0.8249). In both Tet2/3 and Tet tcKO retinas, we observed an increase in the number of PAX6+ amacrine cells (Figures 2E-F; Figure S2; One-way ANOVA followed by a Dunnett’s multiple comparisons test p <0.001; Brown-Forsythe test p = 0.9517). Bipolar cell proportions were also slightly increased in Tet tcKO retinas (Figures 2G-H; Figure S2; One-way ANOVA followed by a Dunnett’s multiple comparisons test p <0.0001; Brown-Forsythe test p = 0.7061).

**Figure 2:**
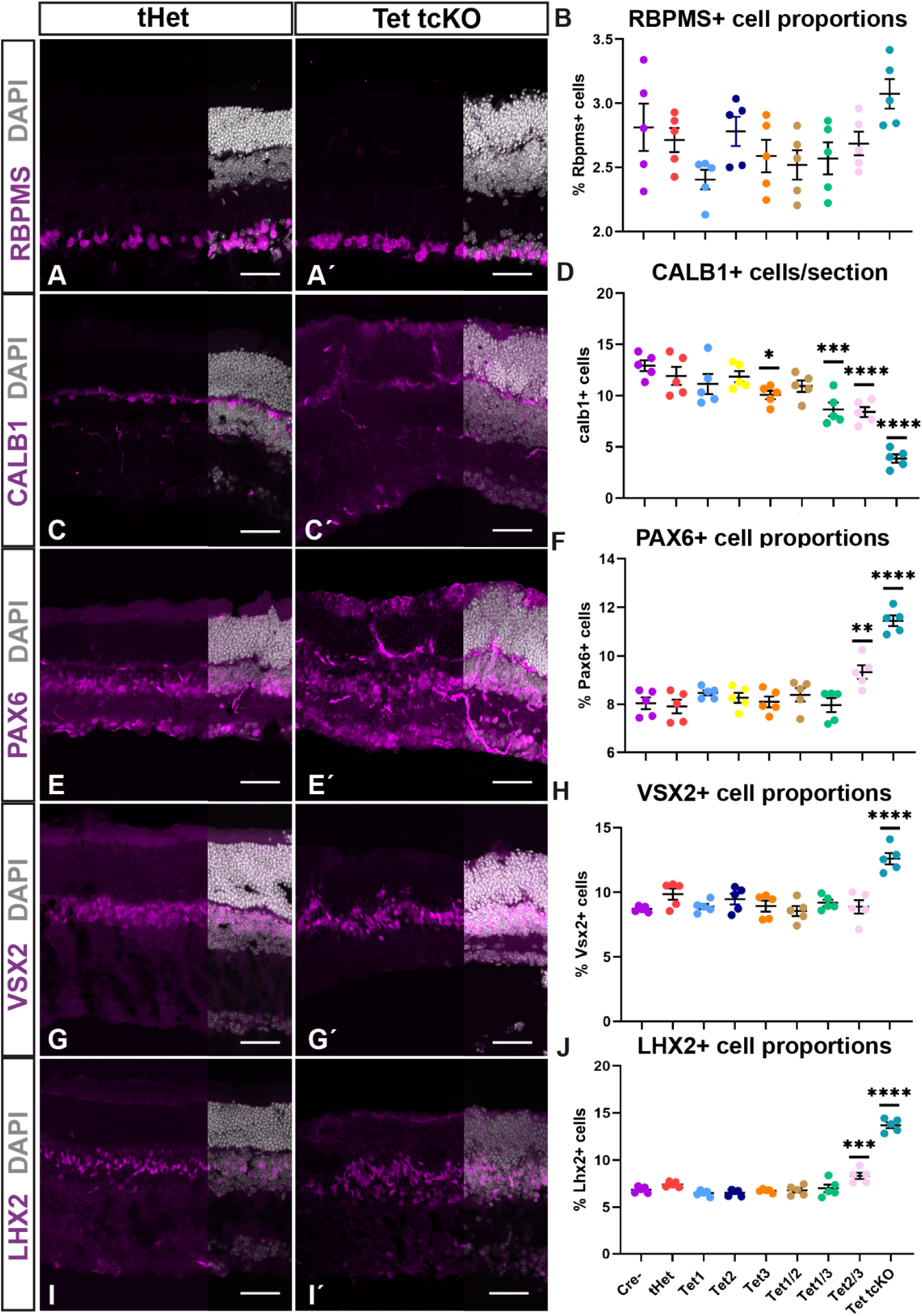
Changes in cell proportions of retinal cell types in TET enzyme conditional mutant retinas. (A, A’) Immunohistochemistry for retinal ganglion cells (RBPMS), (B) Graph showing cell counts of RBPMS+ cell proportions across genotypes. (C, C’) Immunohistochemistry for horizontal cells (CALB1). (D) Graph showing cell counts of CALB1+ cell proportions across genotypes. (E, E’) Immunohistochemistry for amacrine cells (PAX6). (F) Graph showing cell counts of PAX6+ cell proportions across genotypes. (G, G’) Immunohistochemistry for bipolar cells (VSX2). (H) Graph showing cell counts of VSX2+ cell proportions across genotypes. (I, I’) Immunohistochemistry for Müller glia cells (LHX2). (J) Graph showing cell counts of LHX2+ cell proportions across genotypes. Results display the mean + SEM for n=5 for each genotype. Statistics are the result of Ordinary One-Way ANOVA, followed by a Dunnett’s multiple comparisons test compared to Cre- controls. * - p<0.05; ** - p<0.01; *** - p<0.001; **** - p<0.0001.

Recent studies indicate a requirement of the TET enzymes for glial competence of neural stem cells ^89^. We observe an increase in Lhx2+ Müller glia in Tet2/3 cKO and Tet tcKO retinas (Figures 2I-J; Figure S2; Figure S3; One-way ANOVA followed by a Dunnett’s multiple comparisons test p <0.0001; Brown-Forsythe test p = 0.2820). Examination of Müller glia morphology was also assessed through localization of glutamine synthetase (GS), indicating that Müller glia were present, albeit with disorganized morphology in Tet tcKO retinas (Figure S4A). Despite this disorganization, we did not observe glial reactivity as assessed by GFAP in either P21 or 6-week old retinas (Figure S4A). While LHX2 and GS expression suggest specification of Müller glia, we also observed strong colocalization of PAX6, LHX2, and VSX2 within presumptive glial cells (Figure S3). While all of these markers are expressed in mature Müller glial cells ^90–93^, both PAX6 and VSX2 normally display reduced expression in glia compared to that in amacrine and bipolar cells, respectively (Figure S3). However, in Tet tcKO retinas, the prominent co-localization of LHX2, PAX6, and VSX2 is reminiscent of transcription factor expression within RPCs, potentially resulting from a failure to complete glial differentiation and partially accounting for the observed increase in PAX6+ and VSX2+ cells in Tet tcKO retinas. Altogether, we observed subtle yet significant changes in specification of retinal cell fates when TET enzyme expression is lost in the retina.

To determine the potential of cell death influencing retinal cell fate proportions, we assessed the quantity and localization of microglia within the retina. An increase in microglia numbers and localization within the nuclear layers serves as a proxy for microglial activation in response to cell death ^94^. We observe both an increase in microglial number (One-way ANOVA followed by a Dunnett’s multiple comparisons test p <0.0001; Brown-Forsythe test p = 0.3754) and altered laminar distribution of microglia within nuclear (Two-tailed Unpaired t-test p = 0.0018) and plexiform layers (Two-tailed Unpaired t-test p = 0.0079; Figures S4B-E). When examining distribution of microglia within individual nuclear layers, we observe an increase of microglia in all 3 nuclear layers in Tet tcKO retinas compared with tHet controls (Figures S4B-F; ONL: Mann-Whitney test p = 0.0079; INL: Mann-Whitney test p = 0.0476; GCL: Mann-Whitney test p = 0.0079). While this does not rule out the potential for changes in cell fate specification to bias cell type proportions, we suggest that cell death contributes to observed differences in retinal thickness and cell fate proportions in Tet2/3 and Tet tcKO retinas.

Our ERG results indicate the lack of a functional photoreceptor (A-wave) response, potentially through failure of proper photoreceptor differentiation or function in Tet2/3 and Tet tcKO retinas. To further investigate the mechanisms by which photoreceptor dysfunction is occurring, we assessed the expression and localization of known phototransduction related proteins. Consistent with the observed alterations in outer segment development in histological analyses (Figure 1D), we identified attenuated expression and localization of the opsin proteins in both P21 and 6 weeks old retinas (Figures 3A-H). Staining for the short-wavelength cone opsin (OPN1SW) revealed maintained but mislocalized OPN1SW expression (arrows in Figures 3A, A’, C, C’, E, E’, G, G’). While OPN1SW staining was localized to presumptive cone outer segments in Tet tcKO retinas, a significant fraction of OPN1SW protein also mislocalized to the ONL (arrows in Figures 3C, Ć, G, G′). Staining for the rod photopigment Rhodopsin (Figures 3B, D, F, H) showed a near complete loss of Rhodopsin in Tet tcKO retinas (asterisks in Figures 3D, H) that accounts for the reduction in scotopic A-wave observed in this genotype (Figures 1I, L).

**Figure 3:**
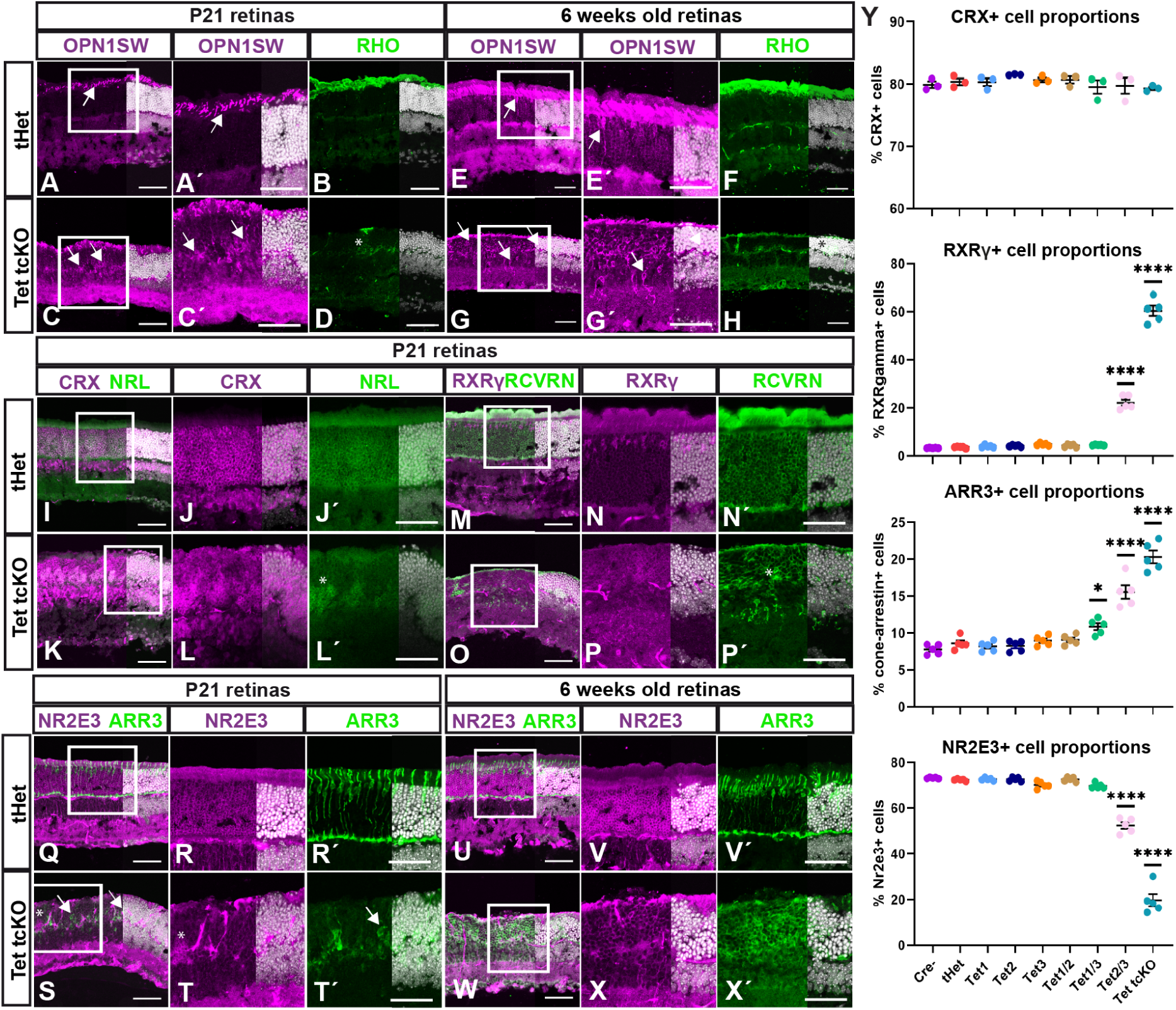
Changes in photoreceptor fate proportions in TET mutant retinas. (A, H’) Immunohistochemistry for S-opsin (OPN1SW) and rhodopsin (RHO) comparing tHet and Tet tcKO retinas at P21 and 6 weeks. (I-L’) Immunohistochemistry for CRX and NRL comparing tHet and Tet tcKO retinas at P21. (M-P’) Immunohistochemistry for RXRγ and RCVRN comparing tHet and Tet tcKO retinas at P21. (Q-X’) Immunohistochemistry for NR2E3 and ARR3 comparing tHet and Tet tcKO retinas at P21 and 6 weeks. (Y) Graphs showing cell counts of CRX+, RXRγ+, ARR3+ and NR2E3+ cell proportions across genotypes. Results display the mean + SEM for n=5 (n=3 in the case of CRX) for each genotype. Statistics are the result of Ordinary One-Way ANOVA, followed by a Dunnett’s multiple comparisons test. *- p<0.05; **** - p<0.0001.

We next performed staining for CRX, a marker of both cone- and rod-photoreceptors, and observed no difference in the proportion of photoreceptors across genotypes (Figures 3I-L;Y; Figures S5A, S5C; One-way ANOVA followed by a Dunnett’s multiple comparisons test p = 0.4774; Brown-Forsythe test p = 0.8572). However, further characterizations of photoreceptor protein expression revealed altered specification of rod versus cone fates. We observed an increase in the proportion of cells that expressed cone transcription factor RXRγ (RXRγ; Figures 3-P’, Y; Figure S5A; One-way ANOVA followed by a Dunnett’s multiple comparisons test p <0.0001; Brown-Forsythe test p = 0.0013) and the cone-arrestin protein (ARR3; Figures 3Q-T’, Y; Figure S5A; One-way ANOVA followed by a Dunnett’s multiple comparisons test p <0.0001; Brown-Forsythe test p = 0.1846) indicative of an increased specification of cone-photoreceptors. This increase in cone-photoreceptors was also observed at 6 weeks (Figures 3U-3X’). Consistent with OPN1SW expression patterns, ARR3 localization was disrupted in Tet mutant retinas (Figure 3; Figure S5A). We observed ARR3 and cone-photoreceptor nuclei mislocalization in deeper layers of the ONL (arrows in Figure 3T’); a pattern maintained in retinas from 6-week Tet tcKO animals (Figures 3W-X’). Conversely, the number of cells expressing master rod-photoreceptor transcription factors NRL ^16–18,94^ (Figures 3I-L’) or NR2E3 ^95–97^ (Figures 3Q-T, Y; Figure S5A) were dramatically reduced in P21 Tet2/3 and Tet tcKO retinas (One-way ANOVA followed by a Dunnett’s multiple comparisons test p <0.0001; Brown-Forsythe test p = 0.0436). Altered photoreceptor fate marker expression was also observed in Tet tcKO retinas at 6-weeks of age (Figure 3F, H, M, N’, O, P’, U, V’, W, X’), the time point used for functional analyses. Our immunohistochemical findings, together with the loss of 5hmC and the enriched expression of TET enzymes in photoreceptor precursors and photoreceptor cells during mouse retinal development (Figure 1B-C) underscore the critical role of TET enzymes and 5hmC in promoting rod-photoreceptor specification. The loss of NRL and NR2E3 but maintained CRX expression suggests that the TET enzymes function upstream of rod fate commitment within specified, CRX+ photoreceptor precursor cells.

Loss-of-function mutations in NRL or NR2E3 result in failure to inhibit cone-photoreceptor fate in presumptive rod-photoreceptors and manifests as enhanced S-cone syndrome in both humans and mice ^16–18,95,97,98^. In our Tet2/3 and Tet tcKO retinas, we observe loss of both NRL and NR2E3 expression, increased proportion of RXRγ+ cones, but a near complete absence of a photoreceptor-mediated A-wave even at the highest light intensities (Figure 1I-L). Photopic B-wave responses, indicative of cone bipolar cell responses, in Tet2/3 and Tet tcKO retinas are detectable, albeit greatly diminished (Figure 1K). To address this discrepancy of increased cone numbers but decreased second order cone-mediated responses, we assessed the presence of synaptic connections and synaptic sublamina using the presynaptic photoreceptor ribbon synapse marker Bassoon ^99^ (CTBP2) or IPL sublamina marker Calretinin ^100^ (CALB2). We observe that Bassoon expression is present but reduced within the inner and outer plexiform layers (IPL and OPL) in both P21 and 6-week old Tet tcKO retinas (Figure S6A). Calretinin staining, which labels the striata of the IPL, is disrupted in both P21 and 6-week old Tet tcKO retinas, with striata becoming undefined in 6-week old Tet tcKO retinas (Figure S6B). Counts of OPL Bassoon+ puncta indicate a significant decrease in the number of ribbon synapses between control and Tet tcKO retinas at P21, with an age-dependent decrease from 3 to 6-weeks (Figure S6C). Our combined results indicate that disruption of retinal cell fate specification, phototransduction protein localization, and alterations to synaptic protein localization combine to cause the reduced visual function in Tet2/3 and Tet tcKO retinas.

### TET enzymes modulate retinal neurogenesis and timing of cone fate specification

During retinal development, photoreceptors are biased to differentiate as a cone ‘default’ state within early retinal development ^98,101^. As development progresses, expression of Prdm1 and Nrl bias photoreceptor precursors to promote rod GRNs, including expression of Nr2e3 ^17,18,96,98,102^. We observed an increase in the proportion of cone-photoreceptors (RXRγ+ cells) at the expense of rod-photoreceptors (NRL+ or NR2E3+ cells) in Tet tcKO retinas. Two plausible scenarios may occur by which enhanced cone-photoreceptor fate is specified in Tet tcKO retinas: 1) TET enzyme loss-of-function results in early cell cycle exit of RPCs during an early ‘competence’ state that promotes cone specification over rod fate; or 2) Photoreceptor fate is initiated properly, but TET enzyme loss-of-function reduces the expression of rod-promoting GRNs through inhibition of NRL and NR2E3 transcription factor expression, thereby adopting a cone fate during periods of rod genesis.

To begin to distinguish between these two scenarios, we performed EdU injections at P0, during the peak of rod genesis, in Cre- and Tet tcKO animals to lineage-trace RPCs (Figure 4). We first utilized a P0 to P1 EdU pulse-chase (Figure 4A) to assess the effect of TET enzyme loss-of-function on cell cycle exit. P0, EdU-labeled RPCs that exited the cell cycle were identified by the lack of co-staining of EdU with VSX2^103–105^. We observed a decrease in the proportion of P0-P1 EdU+ cells that failed to co-express VSX2 in Tet tcKO retinas, indicating reduced neurogenesis and a maintenance of RPC proliferation (Two-tailed Unpaired t-test p <0.0001; Figures 4B-C). However, the total number of VSX2+ RPCs and EdU+ cells in TET tcKO were not statistically different from controls (Two-tailed Unpaired t-test p = 0.3739; Figure 4B; Figure S7A). Additionally, we observed a significant decrease in the proportion (PH3+/DAPI nuclei; Two-tailed Unpaired t-test p = 0.0003) and total number of PH3+ cells (Two-tailed Unpaired t-test p = 0.0009; Figures S7B-D) in Tet tcKO retinas. These data suggest that while numbers of RPCs are properly maintained, the loss of TET enzymes results in decreased mitotic divisions and reduced neurogenic capacity of postnatal RPCs during peak developmental windows of rod fate specification.

**Figure 4.**
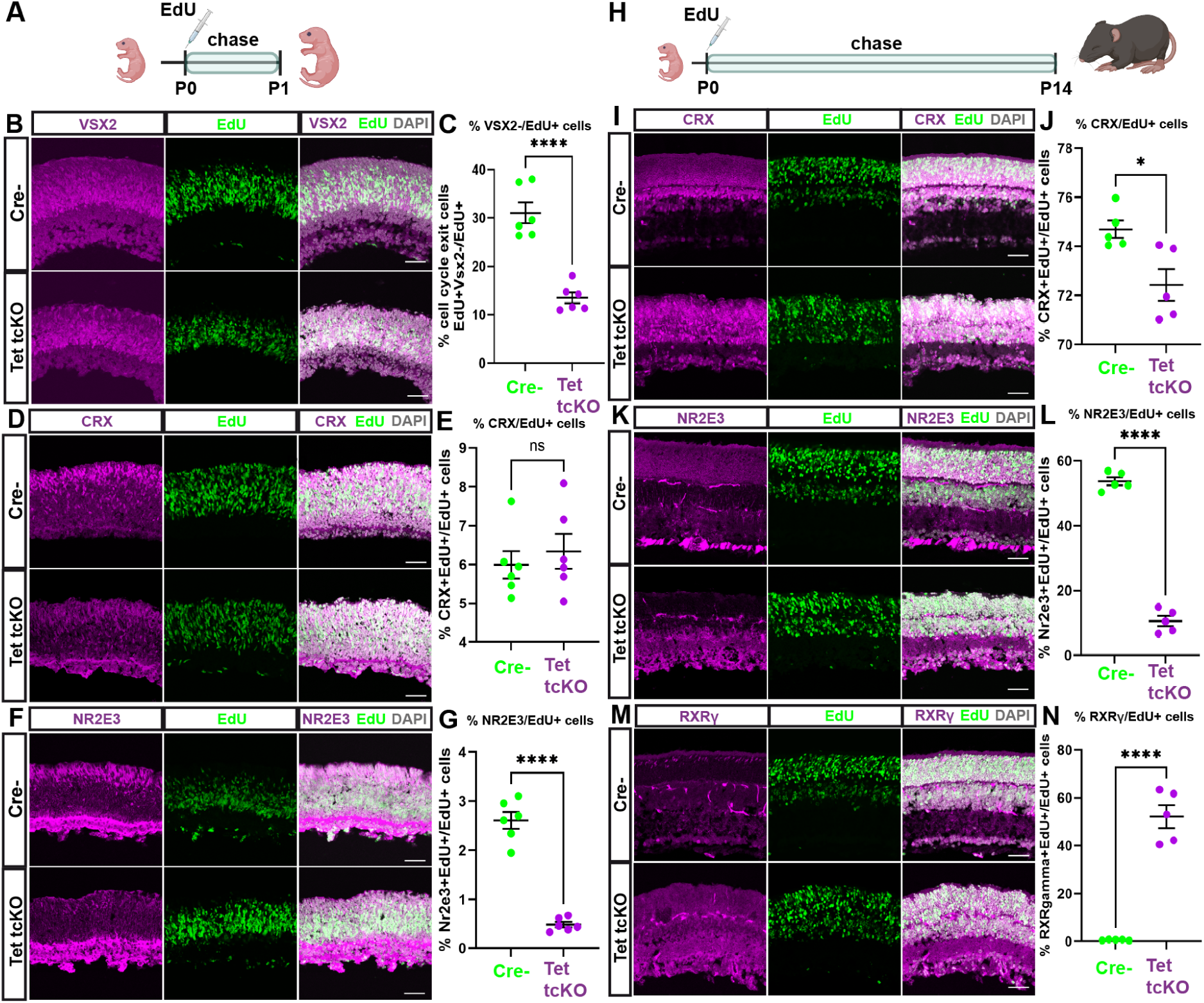
Loss of TET enzymes in RPCs alters retinal neurogenesis and extends the birthwindow of cone-photoreceptors. (A, H) Summary schemas showing the timeline of P0-P1 and P0-P14 EdU experiments. (B, D, F, I, K, M) Immunohistochemistry showing the labeling of progenitor cells (CHX10+), photoreceptors (CRX+), rod-photoreceptors (NR2E3+) and cone-photoreceptors (RXRγ+) in Cre- and Tet1/2/3 cKO retinas at P1 or P14 after EdU injection at P0. (C, E, G, J, L, N) Graphs showing proportion of cells exiting the cell cycle (CHX10+/EdU+), proportions of photoreceptors (CRX+/EdU+), proportions of rod-photoreceptors (NR2E3+/EdU+) and proportions of rod-photoreceptors (RXRγ+/EdU+) Results display the mean + SEM for n=6 (P0-P1) or n=5 (P0-P14) for each genotype. Statistics are the result of a Two-tailed Unpaired t-test; ns: non significant; * - p<0.05, **** - p<0.0001.

Despite observing a decrease in the neurogenic potential of P0-P1 RPCs, co-labeling of EdU+ cells with photoreceptor markers indicate that the numbers of photoreceptors specified during the EdU labeling period were properly maintained (CRX+/EdU+; Figure 4B; Two-tailed Unpaired t-test p = 0.5551). Additionally, the total number of photoreceptors in P1 Tet tcKO mutant and Cre- control retinas were similar (CRX+; Two-tailed Unpaired t-test p = 0.2912; Figures 4D-E; Figure S7E). However, we did observe a reduction in the number of EdU+ cells co-labeled with NR2E3 in Tet tcKO retinas, both as a proportion of total EdU+ cells (NR2E3+/EdU+; Two-tailed Unpaired t-test p <0.0001) and total number of NR2E3+ cells within the Tet tcKO retinas at P1 (Two-tailed Unpaired t-test p = 0.0004; Figures 4F-G; Figure S7F). Altogether, these results indicate that the loss of rod-photoreceptors observed in Tet tcKO retinas occurs at the peak of rod-photoreceptor specification and that postnatal RPCs display reduced neurogenic potential.

We next utilized a P0 pulse-chase to P14 (Figure 4H) to determine the timing of rod- and cone-photoreceptors generation. Using CRX as the common marker of cone- and rod-photoreceptors, we observed a small but significant difference in the number of CRX+ cells generated during the P0-P14 birth window (∼2% change in the proportion of CRX+ cells generated; Two-tailed Unpaired t-test p = 0.0158). Additionally, the total number of CRX+ cells in Tet tcKO retinas compared to controls was also slightly reduced (Two-tailed Unpaired t-test p = 0.0451; Figures 4I-J; Figure S7G). This result is consistent with the reduced number of mitotic divisions and alterations in neurogenesis observed in the P0-P1 EdU experiments. However, when staining for the rod-photoreceptor marker NR2E3, we show a significant decrease in the proportion of NR2E3+ rod-photoreceptors specified from P0 EdU-labeled RPCs (NR2E3+/EdU+; Two-tailed Unpaired t-test p <0.0001), as well as in the total number of NR2E3+ rod-photoreceptors produced in Tet tcKO retinas (Two-tailed Unpaired t-test p <0.0001; Figures 4K-L; Figure S7H).

Consistent with the embryonic specification of cone-photoreceptors in mice ^2^, we observed few P0 EdU labeled RPCs differentiating as RXRγ+ cone-photoreceptors in Cre-control mice (less than 1%). However, EdU+/RXRγ+ co-labeled cells were detected throughout the ONL in Tet tcKO retinas (∼52% of EdU+ cells; Two-tailed Unpaired t-test p <0.0001; Figures 4M-N). We observed a significant increase in the number of total RXRγ+ cones (Two-tailed Unpaired t-test p <0.0001; Figure 4M; Figure S7I), results similar to P21 immunohistological results (Figure 3). Comparisons of NR2E3/EdU double-positive cells in controls with RXRγ/EdU double-positive cells in Tet tcKO retinas reveal that similar proportions of photoreceptors are generated in Tet tcKO and control retinas between P0 and P14 (Figures 4K-N). These findings suggest that the loss of TET enzymes biases immature photoreceptors in the postnatal retina toward adopting a cone rather than a rod-photoreceptor fate.

Altogether, our data favor the hypothesis that TET enzymes function within specified photoreceptors to promote rod fate specification, thereby preventing a cone default state. Deletion of the TET enzymes within RPCs and all resulting progeny prevents rod GRNs from being activated, including loss of NRL and NR2E3 expression. Similar to both NR2E3 and NRL mutant retinas, Tet tcKO retinas display an increase in cone-photoreceptors at the expense of rod-photoreceptors.

### Molecular characterizations of TET enzyme loss-of-function within the mature retina

#### TET enzymes regulate retinal GRNs for determination of photoreceptor fates

Our immunohistochemical characterizations of Tet tcKO retinas led us to hypothesize that TET enzymes are required for promoting rod-photoreceptor fate specification. RNA-sequencing (RNA-seq) experiments on both P21 bulk tissue and isolated single nuclei were performed to further characterize the changes in transcript expression resulting from TET enzyme loss-of-function. In bulk RNAseq experiments, we isolated both RNA and DNA from retinas pooled from two animals for Cre negative (Cre-), Tet1*^loxp/+^*, Tet2*^loxp/+^*, Tet3^loxp/+^ triple heterozygous *Chx10*::Cre-GFP positive (tHet), and Tet tcKO retinas. Three biological replicates were obtained for each pooled condition. RNA was purified for RNA-seq while matched DNA samples were isolated for 5mC and 5hmC profiling (see below; Figure S8A).

Efficiency of our triple conditional knockout approach was assessed by determining the number of reads mapping to TET transcripts, the percentage of TET transcripts overlapping floxed exons, and the canonical splicing efficiency in Tet tcKO retinas compared to controls (Figures S8C-E). While some reads mapped to floxed exons in Tet tcKO samples (Figures S8C-D), we observe a Cre-dependent dosage decrease in the percentage of properly spliced-in reads of Tet floxed exons (spliced in reads over the sum of both spliced in and spliced out reads) across Cre-, tHet, and Tet tcKO retinas (Figure S8E). We also observed a decrease in *TET3* transcript in Tet tcKO retinas while *TET1* and *TET2* transcript expression was not significantly different across genotypes (Figure S8B). The loss of *TET3* transcript expression suggests that deletion of exon4 of *TET3* results in nonsense mediated decay of the mutant transcript.

We observed differential expression of 1102 transcripts in Tet tcKO retinas compared to either Cre- samples (Figure 5A; 627 up-regulated and 475 down-regulated transcripts in Tet tcKO retinas compared with Cre- controls, log2(Fold Change) >1, False Discovery Rate (FDR) < 0.01). 691 (62.70%) transcripts displayed differential expression in both Tet tcKO versus Cre- and Tet tcKO versus tHet pairwise comparisons. All 691 transcripts displaying differential expression in both Tet tcKO pairwise analyses displayed a congruent direction of transcript expression changes across Cre- versus Tet tcKO and tHet versus Tet tcKO pairwise comparisons (331 down-regulated transcripts and 360 up-regulated transcripts).

**Figure 5.**
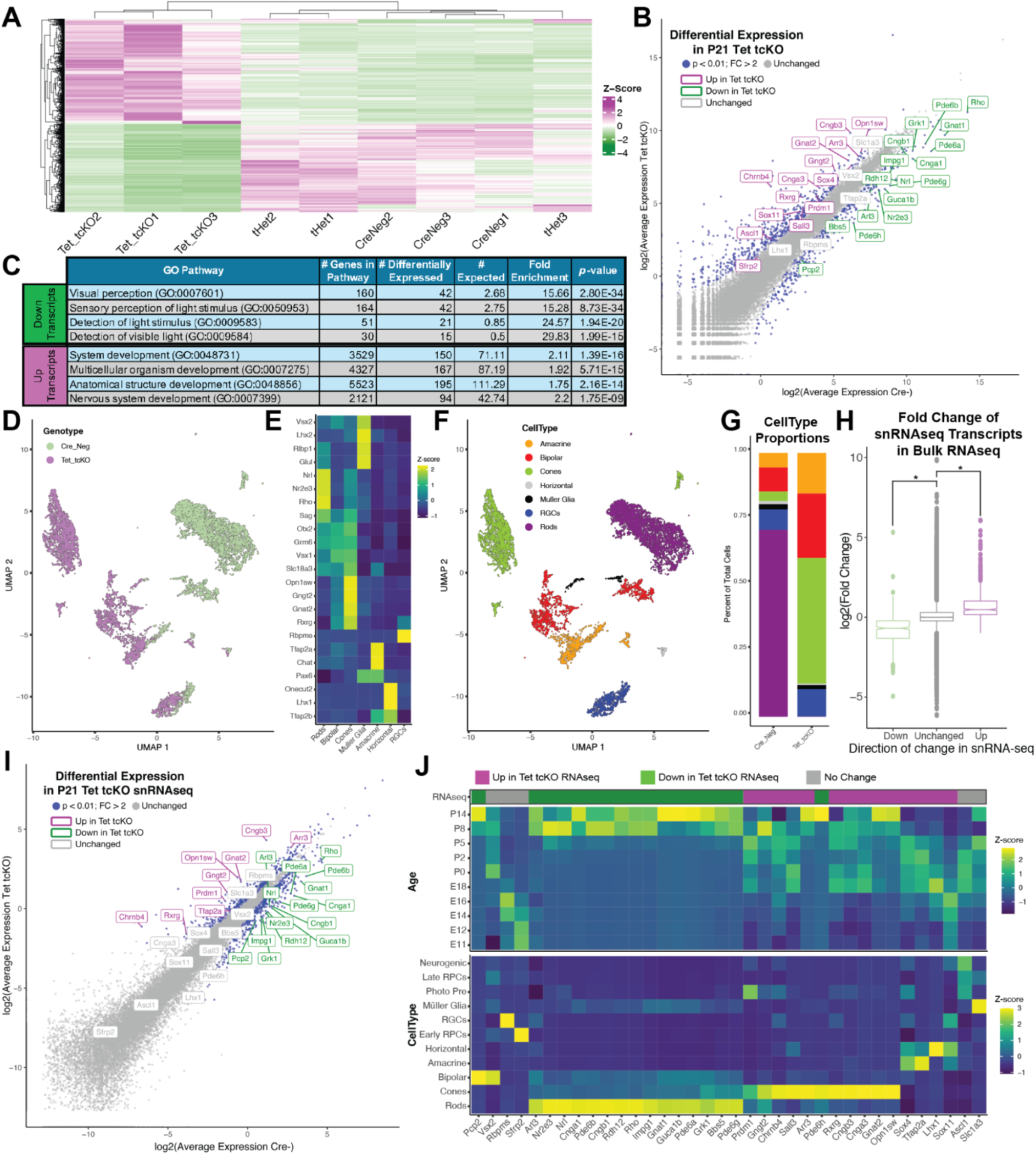
Tet loss-of-function promotes cone-photoreceptors GRNs and loss of rod-photoreceptors. (A) Heatmap of differential transcripts across bulk P21 retinal RNAseq replicates. (B) RNA transcript expression in Cre- and Tet tcKO retinas, colored by differential expression significance and direction of change. (C) Gene Ontology (GO) analysis of differentially expressed transcripts indicating enriched Biological Pathways of Up- and Down-regulated transcripts. (D) UMAP dimension reduction of retinal neurons and glia from snRNAseq on P21 Cre- and Tet tcKO retinas, with cells colored by genotype. (E) Heatmap showing relative expression enrichment of cell type markers within snRNAseq annotated cell types. (F) UMAP dimension reduction of retinal neurons and glia from snRNAseq on P21 Cre- and Tet tcKO retinas colored by annotated cell type. (G) Proportions of annotated cell types by genotype. (H) Boxplots displaying the relative fold change of snRNAseq differentially expressed transcripts in bulk RNAseq experiments. Asterisks represent p-values from Wilcoxon Rank Sum statistical comparisons; * - p < 2.2e-16 (I) Pseudo-bulked, average transcript expression of snRNAseq experiments from Tet tcKO and Cre-. Transcripts are colored by differential expression significance and direction of change. (J) Developmental and cell type expression enrichment of differentially expressed transcripts from Tet tcKO RNAseq experiments in the Clark *et al*. retinal development dataset (PMID: 31128945).

Genes displaying decreased transcript expression include rod-photoreceptor transcription factors, *NRL* (log2 fold change (log2FC)= -2.30; False Discovery Rate (FDR) = 1.03e-15) and *NR2E3* (log2FC = -2.13; FDR = 2.70e-12) and rod phototransduction genes *PDE6A* (log2FC = -3.15; FDR = 1.96e-9), *GNAT1* (log2FC = -2.93; FDR = 1.59e-12), and *RHO* (log2FC = -2.98; FDR = 4.14e-6). Conversely, up-regulated transcripts included cone transcription factor *RXRG* (log2FC = 2.49; FDR = 1.07-e13) and cone phototransduction genes *GNGT2* (log2FC = 2.13; FDR = 3.94e-9)*, GNAT2* (log2FC = 1.94; FDR = 2.63e-14), and *OPN1SW* (log2FC = 1.62; FDR = 1.44e-8). Importantly, we did not observe statistically significant changes in gene expression for photoreceptor transcription factor *CRX,* or cell type-specific markers *RPBMS, TFAP2A, LHX1, VSX2,* or *SLC1A3*, markers of retinal ganglion cells, amacrine cells, horizontal cells, bipolar cells, or Müller glia, respectively (Figure 5B; Table S2). Gene Ontology (GO) pathway analyses of differentially expressed transcripts indicated an enrichment of down-regulated transcripts in biological pathways related to visual perception and detection of light (Figure 5C; Table S3). Conversely, up-regulated transcripts were enriched in pathways related to regulation of development including nervous system development (Figure 5C; Table S3).

Differential expression analysis of Cre- and tHet samples revealed minor differences in RNA transcript abundance across genotypes (37 up-regulated transcripts, 2 down-regulated transcripts, log2(Fold Change) > 1, False Discovery Rate (FDR) < 0.01; Figure S8F; Table S2), indicating minor differences in Cre- and tHet control models. This reduced level of expression changes is consistent with smaller changes in global 5hmC levels observed in the tHet mice.

To further confirm cellular identity changes observed in immunohistochemical data we performed single-nucleus RNA-sequencing (snRNA-seq) of P21 retinas. For both Tet tcKO and Cre- control samples, nuclei from two independent pools of nuclear dissociations (≥2 animals, ≥4 retinas each) were utilized for input into Particle-templated instant partition sequencing (PIPseq)^106^ snRNA-seq reactions. Initial snRNA-seq processing and dimension reduction revealed largely distinct clustering of Tet tcKO nuclei from Cre- controls (Figure 5D). Cell type calls were performed using enrichment of canonical cell type marker genes within clusters (Figures 5E-F)^14^, determining an increase in cone-photoreceptors in Tet tcKOs compared to the Cre- control sample (Figure 5G). Differential expression analysis (Monocle3 fit_models; q-value < 1e-20) across all cells by genotype revealed 518 up- and 523 down-regulated transcripts in Tet tcKO cells compared with Cre- controls (Table S4). Comparisons of differentially expressed transcripts in either snRNA-seq or bulk RNA-seq samples revealed consistent patterns of transcript expression changes between RNA-seq modalities (Figures 5H-I). Examination of the cell type and temporal enrichment of differentially expressed transcripts from Tet tcKO retinas across normal retinal development^14^ confirmed an up-regulation of cone-enriched transcripts and loss of rod transcripts (Figure 5J, bottom). As the peak birth windows of cone-photoreceptors precedes that of rod-photoreceptor ^2,14^, we observed that cone-enriched, up-regulated transcripts in Tet tcKO retinas are normally expressed earlier than down-regulated transcripts during retinal development (Figure 5J, top).

To gain further insight into the cellular enrichment of differentially expressed transcripts from transcript profiling experiments, we examined the cellular expression of transcripts as ‘gene modules’ in the single-cell atlas of the developing mouse ^14^. Consistent with our immunohistochemical data highlighting changes in photoreceptor fate specification, down-regulated transcripts in Tet tcKO retinas display preferential expression within within rod-photoreceptors while up-regulated transcripts are preferentially expressed within cone-photoreceptors (Figure S9). In combination with the immunohistochemical data, both our bulk RNA-seq and snRNA-seq data support a bias in photoreceptor fate specification whereby rod fate is inhibited when the TET enzyme function and 5hmC are lost. Instead, photoreceptor precursors adopt an earlier developmental transcriptional program and are biased to differentiate as a cone default fate.

#### TET enzymes regulate global methylation programs during retinal development

Our immunohistochemical and RNA profiling data highlight a requirement of the TET enzymes for rod-photoreceptor fate specification and differentiation. As the TET enzymes function as regulators of DNA demethylation, we next sought to quantify alterations to DNA methylation patterns that control rod-photoreceptor fate specification and retinal gene expression at base pair resolution.

Previous characterizations of modified cytosines implicated promoter and gene body DNA methylation status as key epigenetic marks for the regulation of transcript expression ^107^. We reanalyzed a temporal series of retinal DNA methylation profiling via Whole Genome Bisulfite Sequencing (WGBS)^27^ to determine the developmental dynamics of DNA methylation status and interrogated how developmental DNA methylation profiles correlate with differentially expressed transcripts in Tet tcKO retinas. As WGBS fails to distinguish between 5mC and 5hmC modifications (Figure S10A), these analyses only compare the levels of modified cytosines (5mC + 5hmC) to unmodified cytosines. We observe that during embryonic and early postnatal development (E14-P0), cone-enriched, up-regulated genes within Tet tcKO retinas display fewer modified cytosines within ± 5000 nucleotides of the transcription start sites (TSS) compared to all other genes (Figures 6A-B; Figure S10B). Conversely, rod-enriched, down-regulated genes in Tet tcKO retinas display more similar to non-differentially expressed genes, levels greater than genes displaying increased expression in Tet tcKO retinas (Figures 6A-B; Figure S10B). These differences in global DNA methylation status are reduced as development progresses (P7-P21; Figures 6A-B; Figure S10B), with the sequence downstream (0 to +5000 base-pairs (bp) of both up- and down-regulated genes displaying less DNA methylation than non-differentially expressed transcripts (Figure 6A; Figure S10B); a result consistent with active maintenance of reduced DNA methylation to facilitate expression of mature photoreceptor genes ^31,86^.

**Figure 6.**
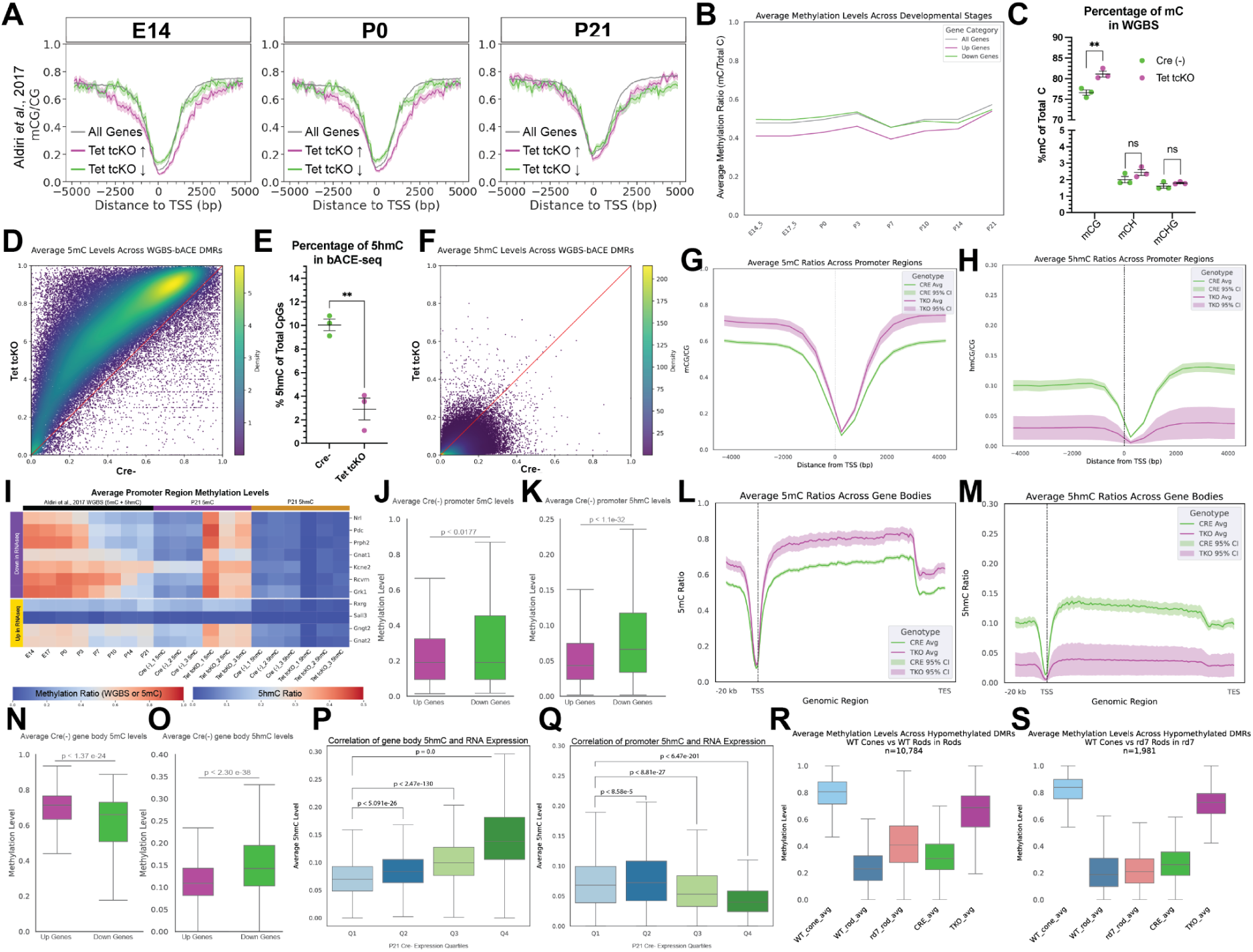
Tet tcKO results in dramatic changes in the retinal methylome. (A) Comparisons of the temporal WGBS methylation patterns at early (E14) mid (P0) and late (P14) timepoints of retinal development across the ± 5kb of the TSS for up- and down-regulated transcripts from Tet tcKO RNAseq experiments. (B) Line graph displaying the average CpG methylation levels across retinal development of the proximal TSS for up- and down-regulated transcripts across retinal development. (C) Scatterplot of the average methylation levels from WGBS performed on P21 Cre- and Tet tcKO samples for mCG, mCH, and mCHG. (D) Density plot of DMRs for 5mC between P21 Tet tcKO and Cre (-) control retinas. (E) Scatterplot of the average 5hmC levels from bACE-seq performed on P21 Cre- and Tet tcKO samples. (F) Density plot of DMRs for 5hmC between P21 Tet tcKO and Cre (-) control retinas. (G-H) Line plot showing the average (G) 5mC or (H) 5hmC levels across the proximal promoter Tet tcKO and Cre (-) control retinas. (I) Heatmaps displaying temporal methylation patterns and 5mC and 5hmC profiles of the promoter regions in Tet tcKO and Cre(-) control retinas for selected, differentially expressed transcripts from RNAseq experiments. (J-K) Boxplots of promoter (J) 5mC and (K) 5hmC levels for genes that display differential transcript expression in RNA-seq experiments (P21 Tet tcKO compared to Cre(-) controls). (L-M) Line plots showing the average (L) 5mC or (M) 5hmC levels across gene bodies in Tet tcKO and Cre (-) control retinas. (N-O) Boxplots displaying average (N) 5mC or (O) 5hmC across gene bodies of differentially expressed transcripts in RNA-seq experiments (P21 Tet tcKO compared to Cre(-) controls). (P-Q) Boxplots displaying average 5hmC levels across the (P) gene body or (Q) promoter for all genes, binned in quartiles by transcript expression levels in P21 Cre(-) RNA-seq. Statistics represent results of a Wilcoxon Rank Sum Test for pairwise comparisons of Quartile 1. (R-S) Boxplots of average methylation profiles for differentially hypomethylated regions identified in comparisons between (R) Rods versus Cones and (S) *rd7* Rods versus Cones in P21 Cre(-) and TET tckO retinal samples and sorted cones, rods, and *rd7* rods. Statistical analyses and G represent the results of unpaired Student’s t-tests (C, G, L-O; ** - p < 0.01, ns - not significant)

We next examined the consequence of Tet enzyme loss-of-function on the retinal methylome. Using the matched DNA from RNA-DNA extractions in RNA-seq experiments, we performed both whole methylome profiling (5mC + 5hmC) via WGBS and 5hmC profiling through bACE-seq ^67,108^ (Figure S10A, C-E). Shallow sequencing of bACE-seq samples in Cre-, tHet, and Tet tcKO retinas indicated that ∼10% of CpG sequences harbor 5hmC modifications within the P21 retina (Figure 1C); a number consistent with 5hmC levels in other profiled nervous system tissues ^109^. Estimates of non-methylated cytosine conversion rates averaged approximately 98% (2% unmodified cytosines failed to be converted; Figure S10C), suggesting robust conversion of cytosine to thymine in bACE-seq reactions.

Deep sequencing of both WGBS and bACE-seq libraries followed by MLML deconvolution analyses ^71^ to resolve 5mC and 5hmC signatures in WGBS provided base-pair resolution of 5mC or 5hmC methylation status (Figures S10D-F). Differentially methylated region (DMR) analyses of WGBS and bACE-seq indicated a large-scale maintenance of cytosine modifications (5mC + 5hmC) in WGBS (191,164 DMRs with up-regulated 5mC+5hmC and 2,672 DMRs losing 5mC+5hmC; Figures 6C-D; Table S5). Our bACE-seq results provide the first base-pair resolution of 5hmC within the mouse retina and identified a dramatic loss of 5hmC across the genome (592,697 DMRs losing 5hmC, 209 DMRs gaining 5hmC; Figures 6E-F; Table S6) in Tet tcKO mutant retinas, consistent with the requirement of the TET enzymes for oxidizing 5mC to 5hmC in post-mitotic cells. Non-CpG methylation levels were lowly abundant and unaffected by deletion of the TET enzymes (Figure 6C). The low levels of non-CpG methylation in the retina are consistent with previous reports ^30^.

Examination of the 5mC levels across the TSS of all genes indicates a global enrichment of 5mC across the proximal promoter region in Tet tcKO retinas (Figure 6G). Conversely, bACE-seq analysis determined that global levels of 5hmC were significantly reduced across the promoter in Tet tckO retinas (Figure 6H). We next sought to understand the biological implications of changes in methylation in the Tet tcKO retinas by comparing the temporal dynamics and cell type specific patterns of methylation across retinal development. Combining the temporal time-series of WGBS methylation profiling ^27^ with our deconvolved 5mC and 5hmC profiling, we next examined how promoters of differentially expressed genes in Tet tcKO retinas are methylated. We observe that promoters of many down-regulated genes in Tet tcKO retinas display high methylation levels in early development that are lost at time points where these genes become highly expressed in the retina (P3-P7 through P21; Figure 6I). Conversely, many up-regulated gene promoters display low levels of DNA methylation that are, on average, less dynamic across development (Figure 6I, bottom). Our P21 methylation profiles observe low levels of 5mC in promoters for all differentially expressed genes in P21 Cre- retinas; however, Tet tcKO retinas display increased 5mC levels at promoter regions, occurring for both up and down-regulated genes (Figure 6I). Similarly, low levels of 5hmC are observed in P21 Cre- samples across both up- and down-regulated genes, with reduced 5hmC levels observed in Tet tcKO knockouts. These results suggest that removing the TET enzymes prevents the DNA demethylation of specific promoters during retinal development, thereby preventing the expression of a subset of genes - those observed as down-regulated in our RNA-seq. We then directly compared promoter and gene body methylation levels of up- and down-regulated genes within the wildtype (Cre-) retina. Promoter 5mC levels between up- and down-regulated transcripts were significantly different despite the mean 5mC levels being similar (Up-regulated gene promoter 5mC = 26.6% of CpGs; Down-regulated promoter 5mC = 28.17%; Figure 6J). Conversely, we observed lower promoter 5hmC levels in genes that are up-regulated compared to down-regulated genes (Figure 6K). Together, these data highlight that up-regulated transcripts in Tet tcKO retinas have less promoter methylation (both 5mC and 5hmC) and therefore do not require DNA demethylation to activate transcription. Conversely, loss of the TET enzymes prevents removal of 5mC and the accumulation of 5hmC modifications at promoters of rod-promoting genes, thereby inhibiting expression of these genes in our mutant model.

Analysis of the methylation across gene bodies followed similar patterns to that of promoter sequences. We observed that gene body 5mC levels are increased across the gene body (TSS to transcription termination site; TTS) compared to the proximal promoter region in Cre- control retinas (Figure 6L). Deletion of the TET enzymes results in an increase of 5mC levels from ∼60% to ∼80% of 5mC-modified CpGs across the gene body (Figure 6L). We also observed a global decrease in 5hmC levels across the gene bodies in Tet tcKO retinas (Figure 6M). We then analyzed the P21 wildtype (Cre-) methylation levels of differentially expressed transcripts from RNA-seq of P21 Tet tcKO retinas to gain a better understanding of how gene body methylation levels affect RNA expression. We observed that, on average, down-regulated transcripts display lower 5mC and higher 5hmC gene body levels in the wildtype retina than up-regulated transcripts (Figures 6N-O), suggesting a requirement of DNA demethylation and potentially 5hmC modifications across the gene body to promote RNA expression. Loss of the TET enzymes maintains gene body 5mC levels of down-regulated genes and inhibits transcription.

Gene body 5hmC levels have previously been associated with higher levels of RNA transcription ^110,111^. To assess the correlation between promoter and gene body 5hmC levels with expression, we binned all genes in RNA-seq experiments by ordered quartiles of expression levels and compared average promoter or gene body 5hmC levels. In agreement with previous reports, we observed that genes with higher levels of RNA transcript expression exhibit higher levels of gene body 5hmC (Figure 6P; Figure S10J). We did not observe a robust correlation of the 5hmC promoter levels with high RNA transcript expression, although highly expressed transcripts did display lower 5hmC promoter levels on average (Figure 6Q; Figure S10K).

To further examine the temporal regulation of methylation and the effect of TET enzyme loss-of-function on the regulation of retinal gene expression, we examined methylation levels of *cis*-regulatory elements (CREs). We utilized a temporal single nucleus Assay for Transposase Accessible Chromatin (snATAC) across retinal development ^112^ to identify all accessible chromatin regions in retinal cells. We then determined the identity of the sites as promoters (± 2000 bp from TSS), gene body, or distal ATAC sequences (ATACseq peaks from Lyu *et al.*, 2021 not associated with promoters or gene bodies). Using the previous characterizations of the temporal dynamics of methylation (5mC + 5hmC)^27^ we identified the CREs that normally undergo DNA demethylation across retinal development (>10% reduction in methylation across development; 42,473 Distal ATAC peaks and 162 promoter sequences). Comparison of our Cre- P21 WGBS methylation patterns in enhancer and promoter sequences correlated well with more mature retina methylation profiles (P10-P21) but poorly with earlier developmental timepoints for both distal ATAC sequences and promoters (Figures S10I-J). This result is consistent with dramatic changes in methylation patterns of accessible regulatory elements across development as retinal cell fates are specified and cells mature. Conversely, the P21 Tet tcKO methylation patterns correlated highest with the E14.5 WGBS methylation patterns for both distal ATAC and promoter sequences (Figure S10H and S10I). Together these results indicate that loss of the TET enzymes within the RPCs early in retinal development promotes an early developmental methylation profile and prevents dynamic changes in methylation patterns.

As we observe an increase in cone-photoreceptor cells at the expense of rod-photoreceptors, we next compared our Cre- and Tet tcKO WGBS methylation ^95,96,113^ profiles to those of sorted rod- and cone-photoreceptors ^30^. As we observe a dramatic decrease in NR2E3 expression in Tet tcKO retinas at both the transcript and protein level, we also compared Tet tcKO methylation profiles to methylation profiles of sorted photoreceptors from *rd7* mice, a model where the rod-promoting NR2E3 transcription factor is mutated and all photoreceptors are cone-like ^30,95,96,113^. Comparisons of methylation levels across hypomethylated regions in rod-photoreceptors or *rd7* rods compared to cones identified that Cre- samples display methylation levels more similar to rods and *rd7* rods than cones (Figures 6R and 6S). However, Tet tcKO methylation profiles are more similar to those of cones in both comparisons, indicating maintained methylation of genomic regions in Tet tcKO retinas that normally undergo demethylation in rods (Figures 6R-S). Importantly, loss of NR2E3 in the *rd7* model displays a methylation pattern more similar to rods than cones, suggesting that methylation patterns in rods are established upstream of NR2E3 function (Figure 6R and 6S). Examination of methylation status of genomic regions that are hypomethylated in cones, however, indicates that Tet tcKO retinas maintain high methylation of these sites (Figures S10K-L), suggesting that while Tet tcKO retinas display expression of numerous cone marker genes, active DNA demethylation is required to fully establish the proper cone methylome. While Tet enzyme loss of function promotes cone-photoreceptor fate, cone-photoreceptor function is likely impaired due to incomplete maturation of the cells as a consequence of altered DNA methylation.

Association of 5hmC DMRs with genomic features indicate that >30% of 5mC and 5hmC DMRs were localized to Distal Intergenic regions (Figure S10M). Distal enhancer sequences displayed the highest average change in 5hmC modifications in Tet tcKO retinas (Figure S10N). As methylation status affects transcription factor binding to DNA ^114^, future analysis into the significance of 5hmC deposition on enhancer sequences will determine the positive or negative effects of 5hmC on enhancer activity. However, examination of the distribution of DMRs across snATAC peaks in photoreceptor gene loci provided further support for the requirement of DNA demethylation for rod development (Figure S11). Cone-photoreceptor loci (*Rxrg*, *Arr3*, and *Opn1sw*) showed little change in 5mC levels (purple bars) across accessible peaks (blue bars). 5hmC levels displayed larger changes with loss of the TET enzymes. Conversely, in both genetic loci of photoreceptor precursor cells (*Crx, Otx2,* and *Prdm1*) and rod-photoreceptors (*Nrl, Nr2e3,* and *Rho*), we observed large regions of DMRs that overlapped regulatory sequences that gained 5mC and lost 5hmC (Figure S11). Further insight into the mechanisms by which the Tet enzymes are targeted to these loci are required.

### Loss of 5hmC and TET3 expression in human retinoblastoma

Recent evidence implicates altered TET enzyme expression (mRNA and/or protein) within multiple cancers. For example, TET2 loss-of-function mutations are frequently observed in lymphoid and myeloid cancers ^115^ and gliomas frequently exhibit large reductions in 5hmC levels, reduced TET expression, and reduced TET nuclear localization ^116^. In the retina, the primary intraocular cancer in pediatric patients is retinoblastoma, occurring through either mutations in the RB1 gene or MYCN amplification ^117^. Interestingly, maturing cone-photoreceptor cells represent the retinoblastoma cell type of origin ^118–120^. As our Tet tcKO lose TET enzyme expression, display globally reduced levels of 5hmC, and display gene expression and methylation patterns of immature cones, we next examined similarities between human retinoblastoma samples and TET tcKO mutants.

Sections from formalin-fixed, paraffin-embedded (FFPE) retinoblastoma cases of enucleation were stained for 5hmC and TET3, showing loss of both 5hmC modifications and TET3 nuclear expression within the retinoblastoma tumors compared with surrounding, unaffected tissue (Figures 7A-G). To determine the extent that Tet tcKO retinas recapitulate features of retinoblastoma, we performed RNA-seq on retinoblastoma samples and normal retina controls. Differential expression between retinoblastoma tumor and control, healthy retina observed large-scale changes in the transcriptome (3152 up-regulated and 3718 down-regulated transcripts, Table S7). Down-regulated transcripts included genes pivotal to rod-photoreceptor function including RHO, PDE6A, and SAG (Figure S12A). GO pathway analysis highlighted the depletion of transcripts in pathways for synaptic transmission, neural development, and sensory perception of light (Figure S12B). Conversely, transcripts displaying enrichment in retinoblastoma tumor samples were associated with DNA replication and cell division (Figure S12C), consistent with ongoing proliferation within the tumor.

**Figure 7.**
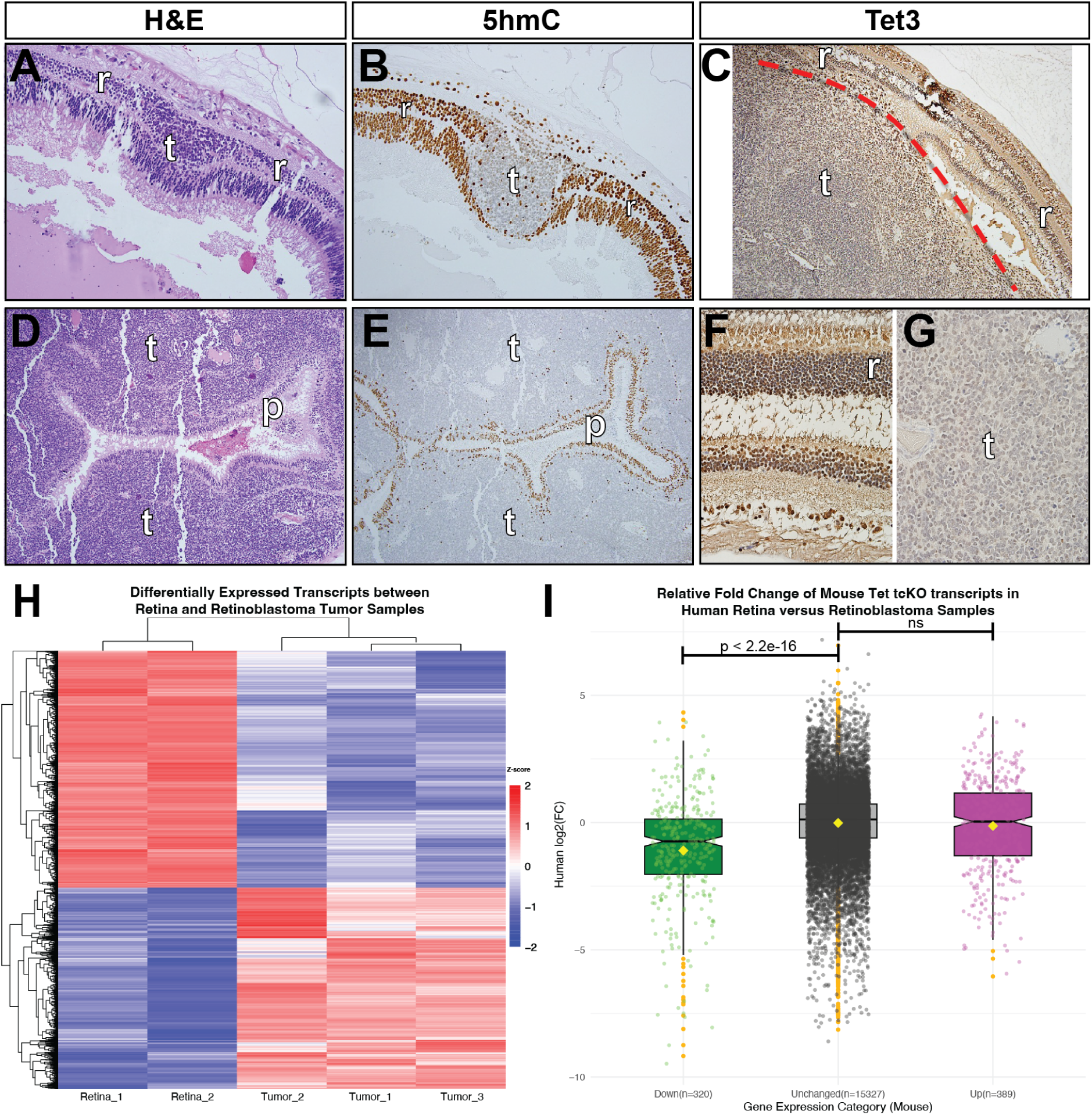
TET3 and 5hmC are lost in Retinoblastoma coincident with large changes in gene expression. (A and D) H&E staining of (A) retinoblastoma tumor (t) surrounded by normal retinal tissue (r) or (D) a large retinoblastoma tumor surrounding few remaining photoreceptors (p). (B and E) Immunocytochemistry localizing 5hmC within the retina (r) and photoreceptors (p), with absence of 5hmC within tumor cells. (C, F and G) Tet3 immunocytochemistry of human retina (r) and retinoblastoma tumor cells (t). (F) Tet3 is localized to nuclei throughout the retinal layers, but (G) Tet3 is weakly detected in tumor cells. (H) Heatmap of differentially expressed transcripts from RNAseq experiments of normal human retinal tissue and retinoblastoma tumors across individual replicate samples. (H) Boxplots showing the relative fold change in human retina versus human retinoblastoma RNAseq of the human orthologs of mouse differentially expressed transcripts between P21 Tet tcKO and Cre-retinal samples. (I) Boxplots displaying the relative fold-change of differentially expressed orthologs of the Tet tcKO differentially expressed transcripts within the human retinoblastoma RNAseq comparisons. Genes are categorized by direction of change in Tet tcKO RNAseq, with outlier values highlighted by orange dots. Notch within boxplots represents the median value, with the yellow diamond representing the category mean. Statistics represent the results of pairwise Wilcoxon Rank Sum analyses in comparison to unchanged control transcripts.

We next examined the extent to which orthologs of differentially expressed genes from Tet tcKO retinas were similarly dysregulated in human retinoblastoma samples. We observe that down-regulated transcripts in Tet tcKO samples were on average, similarly decreased in retinoblastoma samples. These results reflect the loss of rod-photoreceptors in Tet tcKO retinas and cone origin, and subsequent rod depletion, in retinoblastoma samples. However, depleted transcripts in human retinoblastoma samples compared to normal retina were not likewise down-regulated in Tet tcKO retinas (Figure S12D). We then conducted the complementary analyses to determine gene expression signatures resulting from altered DNA methylation in TET tcKO retinas were similarly changed in retinoblastoma. Tet tcKO up-regulated transcripts, however, were not similarly increased in human retinoblastoma, displaying no difference in comparison to the unchanged control genes and reflecting the large proliferative signature of the retinoblastoma tumor ^119^.

To further compare rod versus cone gene expression changes in Tet tcKO and human retinoblastoma samples, we identified a set of genes displaying enrichment in either cone- or rod-photoreceptors using a temporal time series of scRNA-seq performed on human retinal development (Table S8)^83^. In both Tet tcKO and retinoblastoma samples, cone-enriched transcripts displayed increased expression compared to control samples, while rod-enriched transcripts were depleted (Figures S12E-F). Our results highlight that 5hmC and TET3 expression are likewise lost in retinoblastoma as in our Tet tcKO retinas. Similarly, Tet tcKO up-regulated and cone-enriched transcripts display increased expression in human retinoblastoma samples. However, the extent to which loss of TET3 and 5hmC is causative to or a direct consequence of retinoblastoma remains to be elucidated.

Altogether, our results highlight the requirement of the TET enzymes and DNA demethylation for the proper specification of rod-photoreceptors during retinal development. Our results highlight novel mechanisms regulating retinal cell fate specification, placing the TET enzymes upstream of rod fate choice. We show that expression of both NRL and NR2E3, master rod-photoreceptor transcription factors, are inhibited when DNA demethylation is impaired. Base-pair resolution profiling of both 5mC and 5hmC highlight the large-scale dynamics of TET-mediated DNA demethylation, including maintenance of 5mC modifications within the NRL and NR2E3 loci in Tet tcKO retinas. Therefore, we hypothesize a model by which cone-photoreceptor fate is promoted during early development because cone-promoting genes are not actively regulated by DNA demethylation. Rod-photoreceptor fate, however, requires TET-mediated DNA demethylation within post-mitotic photoreceptor precursors, whereby expression of NRL and NR2E3 are induced to inhibit cone fate, enabling the developmental switch from cone to rod fate specification (Figure 8).

**Figure 8.**
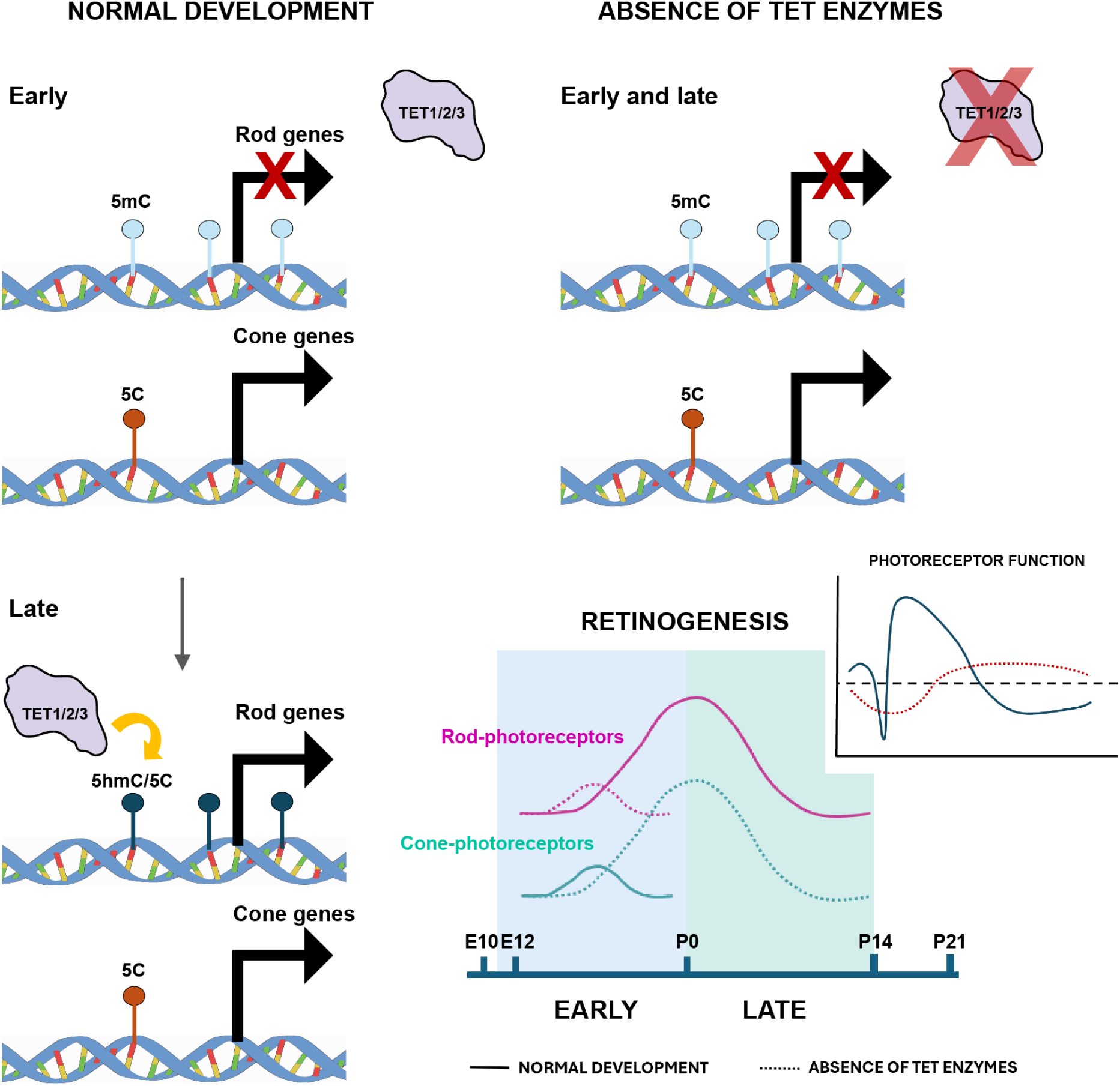
Model of the molecular mechanisms by which TET-mediated DNA demethylation regulates photoreceptor fate decisions. During early retinal development rod-photoreceptor genes (including NRL and NR2E3) are methylated. As cone-photoreceptor genes are lowly methylated, cone fate is favored. As development progresses, the TET enzymes mediate demethylation of rod-promoting genes. In the absence of TET enzymes, demethylation of NRL, NR2E3, and other genes is inhibited, preventing expression and leading to a retina where all photoreceptors are specified as cones.

## DISCUSSION

Here we describe a novel mechanism for the regulation of photoreceptor fate choice within the developing retina. Using a conditional knockout mouse model to remove the TET enzymes from developing RPCs, we show that TET enzyme-mediated DNA demethylation is required for specification of rod-photoreceptors. Removal of the TET enzymes results in a cone-rich retina, while proportions of other major retinal cell types are largely maintained. We observe a requirement of the TET enzymes within CRX+ photoreceptor precursors for the expression of master rod-photoreceptor transcription factors NRL and NR2E3. Our results, including comprehensive transcriptional profiling in conjunction with base-pair resolution profiling of 5hmC and 5mC modifications indicate that oxidation of 5mC to 5hmC is vital to retinal development and function. We observed that 5hmC is maintained across ∼10% of all CpGs and that 5hmC modifications over gene bodies is correlated with gene expression levels. Furthermore, profiling of human retinoblastoma samples indicates an attenuation to TET3 expression and 5hmC deposition levels, suggesting that DNA demethylation and 5hmC modifications may contribute to the pathogenesis of the disease.

Previous studies have identified that numerous promoters and gene bodies of photoreceptor genes (including NRL and NR2E3) are methylated in progenitors but exhibit low DNA methylation levels and high chromatin accessibility in mature photoreceptors ^29–31^. Combined with the enriched expression of demethylation pathway components within the photoreceptor lineage (Figure 1B), this has led to the hypothesis that DNA demethylation, and therefore the TET enzymes, are vital for photoreceptor fate determination, maturation, and function. Previous studies in Tet2^-/-^;Tet3^-/-^ zebrafish show retinal ganglion cell differentiation deficiencies including altered optic nerve development ^61^. However, in our studies we did not find significant differences in the number of retinal ganglion cells. A recent similar study observes that conditional deletion of the TET enzymes results in a thinner optic nerve and progressive retinal degeneration ^86^, suggesting that prolonged dysregulation of DNA demethylation results in cell death. This result is supported in our model by the increase in nuclear-localized microglia in TET tcKO retinas (Figures S4B-F). Zebrafish Tet2/3 mutant retinas display problems in the terminal differentiation of photoreceptors and the absence of their outer segments ^61,121^. However, in our studies, the absence of TET enzymes leads to an increase in cone-photoreceptors at the expense of rod-photoreceptors. This discrepancy in models may be the result of either species-specific differences in cell type composition or compensation from Tet1 as Tet enzymes show partial functional redundancy ^56,122,123^. In our Tet tcKO mice, we also observe photoreceptor dysfunction and reduced transmission of visual signals to second-order neurons (i.e. bipolar cells). Lineage tracing analyses in Tet tcKO retinas highlight the normal specification of photoreceptor numbers, but altered photoreceptor subtype fate specification. Histological and RNA-seq analyses (Figures 2-5) confirmed both lack of NRL and NR2E3 RNA transcript and protein expression, leading to enhanced specification of a cone-photoreceptor ‘default’ state.

Previous studies have shown that loss of either NRL or NR2E3 leads to development of cone-dominant retinas that display decreased expression of rod-specific genes such as Rhodopsin, Recoverin or GNAT1 and increased expression of cone-specific genes including PDE6C, GNAT2 or OPN1SW. In both NRL and NR2E3 mutant mouse models, failure to fully specify rod-photoreceptors leads to retinal degenerations. In humans, disruption of either NRL or NR2E3 expression causes enhanced S-cone syndrome, characterized by supranormal blue cone function due to an increased proportion of S-cones and night blindness due to the absence of rod-photoreceptors ^95,97^. This phenotype has been also described in NR2E3-null human organoids, which show the disruption of photoreceptor cell fate and maturation ^124^. Expression profiling of the Tet tcKO retinas indicates similar changes in gene expression to NRL or NR2E3 loss-of-function models, leading us to propose a model whereby TET enzymes are upstream of NRL and NR2E3 for specification of rod-photoreceptors within post-mitotic photoreceptor precursor cells (Figure 8).

Our methylation profiling experiments (Figure 6) highlight global changes in the retinal methylome in Tet tcKO retinas. We observe the loss of 5hmC and maintained 5mC in both the NRL and NR2E3 gene loci (Figure S11), indicating that the TET enzymes likely regulate NRL and NR2E3 expression and rod-photoreceptor GRNs. Therefore, NRL and NR2E3 deficient photoreceptor precursors adopt a cone default fate in Tet tcKO retinas, highlighted by expression of the cone transcription factor RXRγ. However, despite the increase in the number of cone-photoreceptors, the Tet tcKO cones do not display DNA methylation profiles similar to those of mature cones. We suggest that the TET enzymes are required for full maturation of cone photoreceptors and that loss of the Tet enzymes inhibits cone function and alters retinal morphology. These results are consistent with additional Tet mutant mouse and zebrafish models recently reported ^86,121^.

Our results highlight the requirement of active DNA demethylation within photoreceptor precursors to promote rod-photoreceptor fate specification. CRX, the cone-rod homeobox transcription factor, acts upstream of both NRL and NR2E3. CRX initiates expression in mice at E12.5 in post-mitotic specified photorector precursors ^125–127^. CRX binding coordinates the expression of rod- or cone-photoreceptor genes required for individual photoreceptor subtype specification ^128,129^. Mutations in the CRX sequence or changes in its binding sites lead to several retinal diseases including Leber Congenital Amaurosis (LCA), Cone-rod Dystrophies (CoRD), and Retinitis Pigmentosa (RP) ^65,130,131^, as well as changes in chromatin remodeling in specific target sites ^129^. We observe that CRX expression is unchanged in Tet tcKO retinas despite alterations to the methylation patterns in the *Crx* locus (Figure S11), suggesting that Tet tcKO retinas have the potential to generate rods. However, recent work has explored the consequence of DNA methylation on CRX binding to DNA, identifying altered CRX binding to the CRX consensus motif (TAATCC) by the presence of cytosine methylation modifications and DNA demethylation intermediates (5mC, 5hmC, 5fC and 5caC)^132^. Our methylation profiling results highlight that cone-enriched, up-regulated genes in Tet tcKO retinas display reduced 5mC levels and less temporal dynamics across development (Figure 6, Figure S11). Conversely, the loss of TET enzyme expression prevents the demethylation of rod-enriched transcripts. Therefore we suggest that the cone-enriched Tet tcKO retina may be a result of failure of CRX binding to rod-enriched gene loci because of altered DNA methylation.

While we have identified plausible mechanisms for why cone fate is promoted in Tet tcKO retinas, we do not yet fully understand how the TET enzymes are targeted to specific rod-promoting gene loci to promote DNA demethylation within photoreceptor precursors or how this process is regulated in a temporal manner consistent with the temporal specification of retinal cell fates. The specific targeting of the TET enzymes to DNA sequences is a topic of widespread interest. Although TET enzymes have partially redundant functions in central nervous system development, recent evidence suggests only slight sequence preference for each TET enzyme ^133^. It is more likely that the TET proteins interact with other cofactors to drive sequence and context-specific DNA demethylation. The GADD45 proteins are reported to promote demethylation by mediating TET enzyme localization to DNA ^134–138;^ however, the significance of the GADD45 proteins for controlling

DNA demethylation remains controversial ^139–141^. The potential role of the GADD45 proteins in regulating temporal DNA demethylation patterns is intriguing given that *GADD45A* and *GADD45G* display enriched expression specifically in early and late neurogenic cells, respectively ^14,83^. Alternatively, LIN28A, an RNA binding protein involved in the control of RPC proliferation and neurogliogenesis ^142^ is known to recruit TET1 to DNA to promote DNA demethylation ^143,144^. Furthermore, the CXXC proteins interact with the TET enzymes to maintain and stabilize the demethylated state of specific DNA loci ^145^. CXXC4/CXXC5 are both expressed within the developing retina, however the function of these proteins within the retina remains unknown. Further studies will focus on deepening our understanding of the molecular mechanisms that DNA methylation marks are recognized and targeted for site-specific demethylation in control of retinal development.

## DATA AVAILABILITY

Raw sequencing data and processed files for RNAseq, snRNAseq, WGBS and bACEseq experiments are available through GEO under accession numbers GSE288096, GSE288097, GSE288098, and GSE288100.

## ACKNOWLEDGEMENTS

This work was supported by National Eye Institute of the National Institutes of Health grants R00EY027844 and R01EY035381 (BSC), R01EY012543 (SC), R01EY036368 (PAR), T32EY013360 (CH), K08EY026654 (RCR), R01EY030989 (RCR) and by P30EY002687 (WashU Department of Ophthalmology and Visual Sciences) and P30EY007003 (University of Michigan Vision Research Center) Core Services Grants. Research to Prevent Blindness supported BSC, PAR, and RCR with individual Career Development Awards and unrestricted funds to either the WashU Department of Ophthalmology and Visual Sciences or the Kellogg Eye Center at the University of Michigan. RCR was also supported by the A. Alfred Taubman Medical Research Institute, the Beatrice and Reymont Paul Foundation, March Hoops to Beat Blindness, and Leonard G. Miller Endowed Professorship and Ophthalmic Research Fund at the Kellogg Eye Center. Additional support for this research was provided by Grossman, Elaine Sandman, Marek and Maria Spatz (endowed fund), Greenspon, Dunn, Avers, Boustikakis, Sweiden, and Terauchi research funds. The. The NCI Cancer Center Support Grant also supported this work (P30CA046592). The authors thank the Genome Technology Access Center at the McDonnell Genome Institute at Washington University School of Medicine for help with genomic analysis. The Center is partially supported by NCI Cancer Center Support Grant (P30CA91842) to the Siteman Cancer Center and by ICTS/CTSA (UL1TR002345) from the National Center for Research Resources (NCRR).

## AUTHOR CONTRIBUTIONS

BSC supervised the project. BSC, IHN, AU, SC, PAR, and JRE designed experiments. IHN, AU, XZ, WJ, CH, EH, LQ, FM, CA, PAR, JRE, and BSC performed experiments and interpreted the data. MMD provided the TET conditional alleles and provided insights into experimental design. RR, AC, QL, and FM performed immunohistochemical and RNA-seq experiments on primary retinoblastoma tissue. IHN and BSC wrote the manuscript with input and editing performed by all authors.

## Notes

### Competing Interest Statement

The authors have declared no competing interest.

