## Supplementary material for "Active DNA demethylation is upstream of rod-photoreceptor fate determination and required for retinal development": Figure S1-S12

### SUPPLEMENTAL DATA

### SUPPLEMENTAL FIGURES

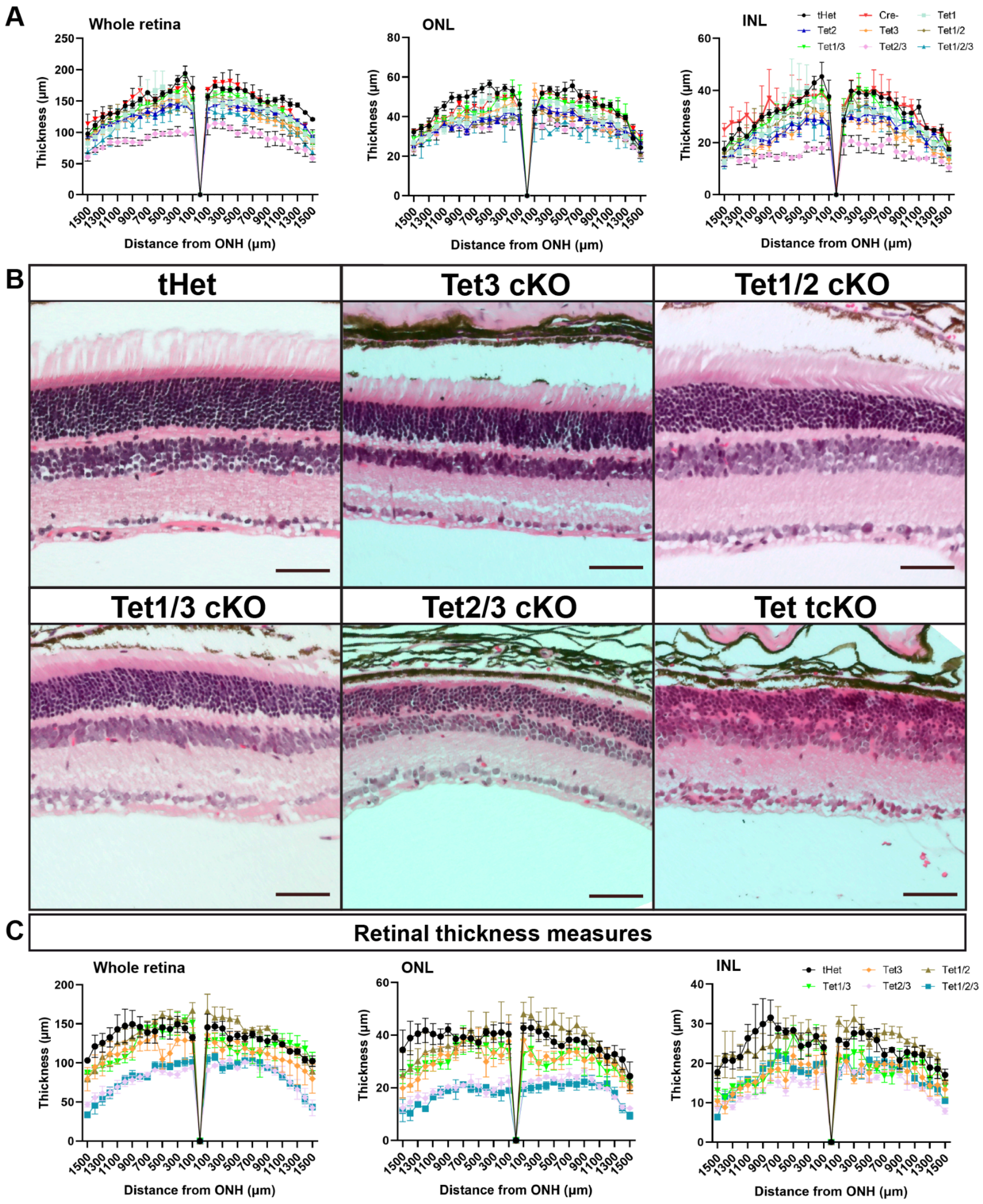


**Supplemental Figure S1. Characterization of the morphological changes driven by TET enzyme loss-of-function. Related to Figure 1.** (A) Whole retina, outer nuclear layer (ONL) and inner nuclear layer (INL) thickness measures at different eccentricities from the optic nerve head (ONH) in P21 retinas. Results display the mean + SEM for n=3 for each genotype. (B) H&E staining of an allelic series of TET conditional P21 mutants. (C) Whole retina, outer nuclear layer (ONL) and inner nuclear layer (INL) thickness measures at different eccentricities from the optic nerve head (ONH) in 6 weeks old retinas. Results display the mean + SEM for n=3 for each genotype.

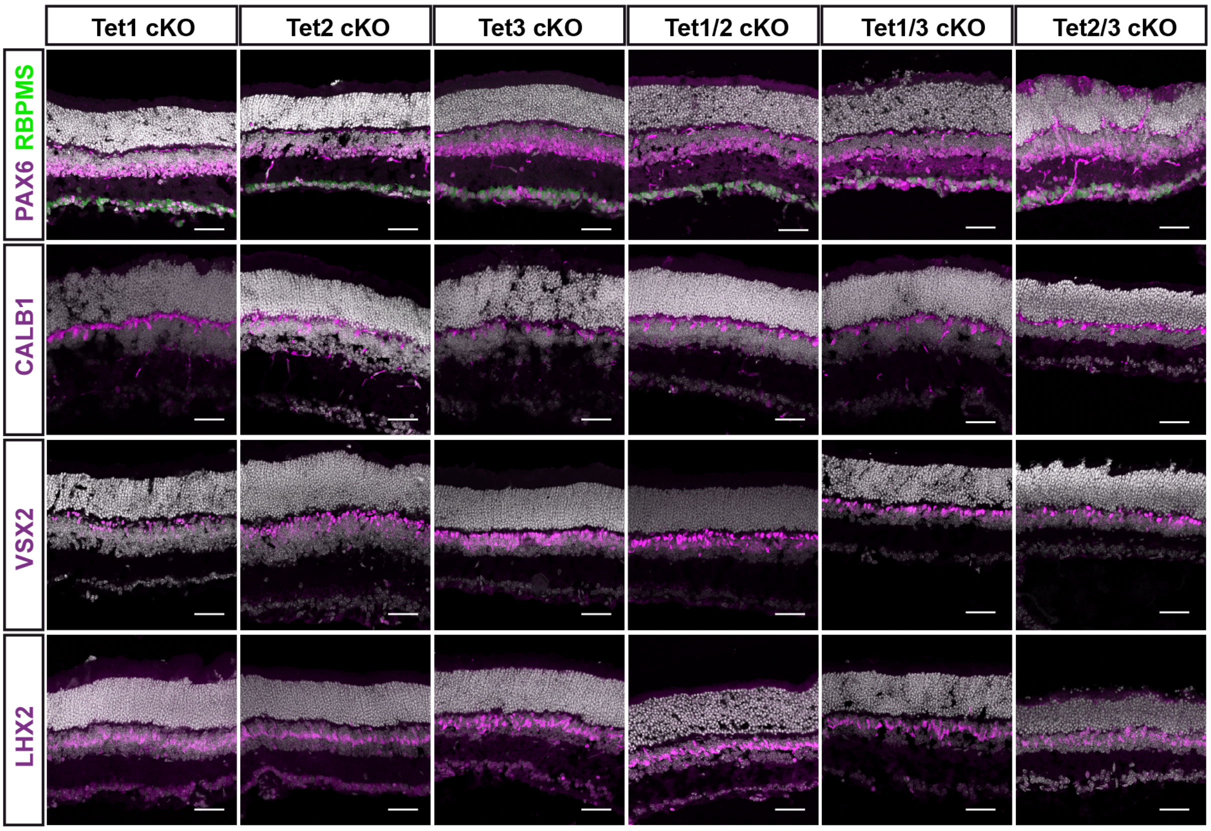


**Supplemental Figure S2. Immunohistochemical characterizations of cell types across the TET enzyme allelic series of conditional mutants. Related to Figure 2**. Immunohistochemistry for retinal ganglion cells (RBPMS), amacrine cells (PAX6); horizontal cells (CB); bipolar cells (VSX2); Müller glia cells (LHX2) markers that show changes in cell proportions of some retinal cells types when TET enzymes are absent.

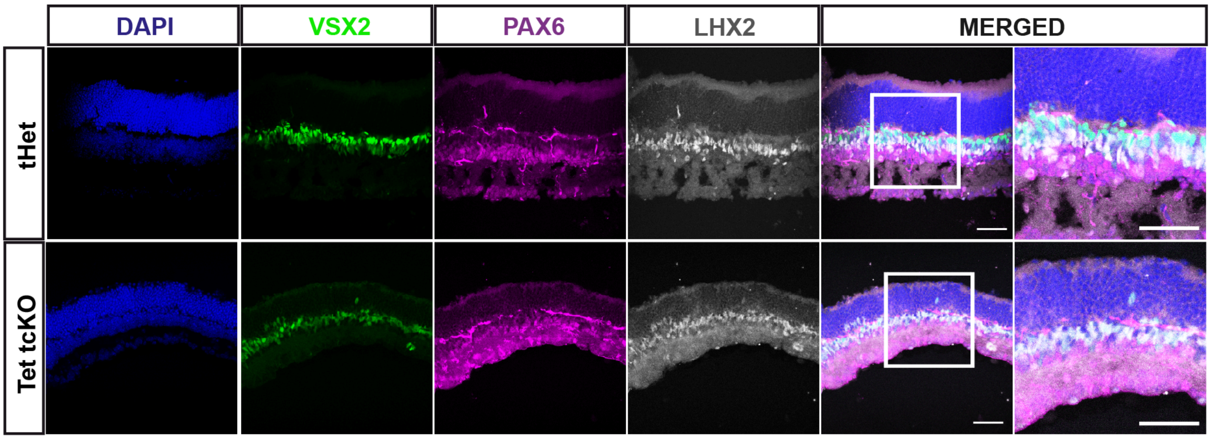


**Supplemental Figure S3. Loss of TET enzymes results in high levels of RPC transcription factors co-expression within presumptive Müller glia. Related to Figure 2 I-J.** Immunohistochemistry for bipolar cells (VSX2); amacrine cells (PAX6) and Müller glia cells (LHX2) markers in tHet and Tet tcKO retinas that show prominent co-localization of VSX2, PAX6, and LHX2 and in presumptive Müller glia in Tet1/2/3 cKO retinas when TET enzymes are absent.

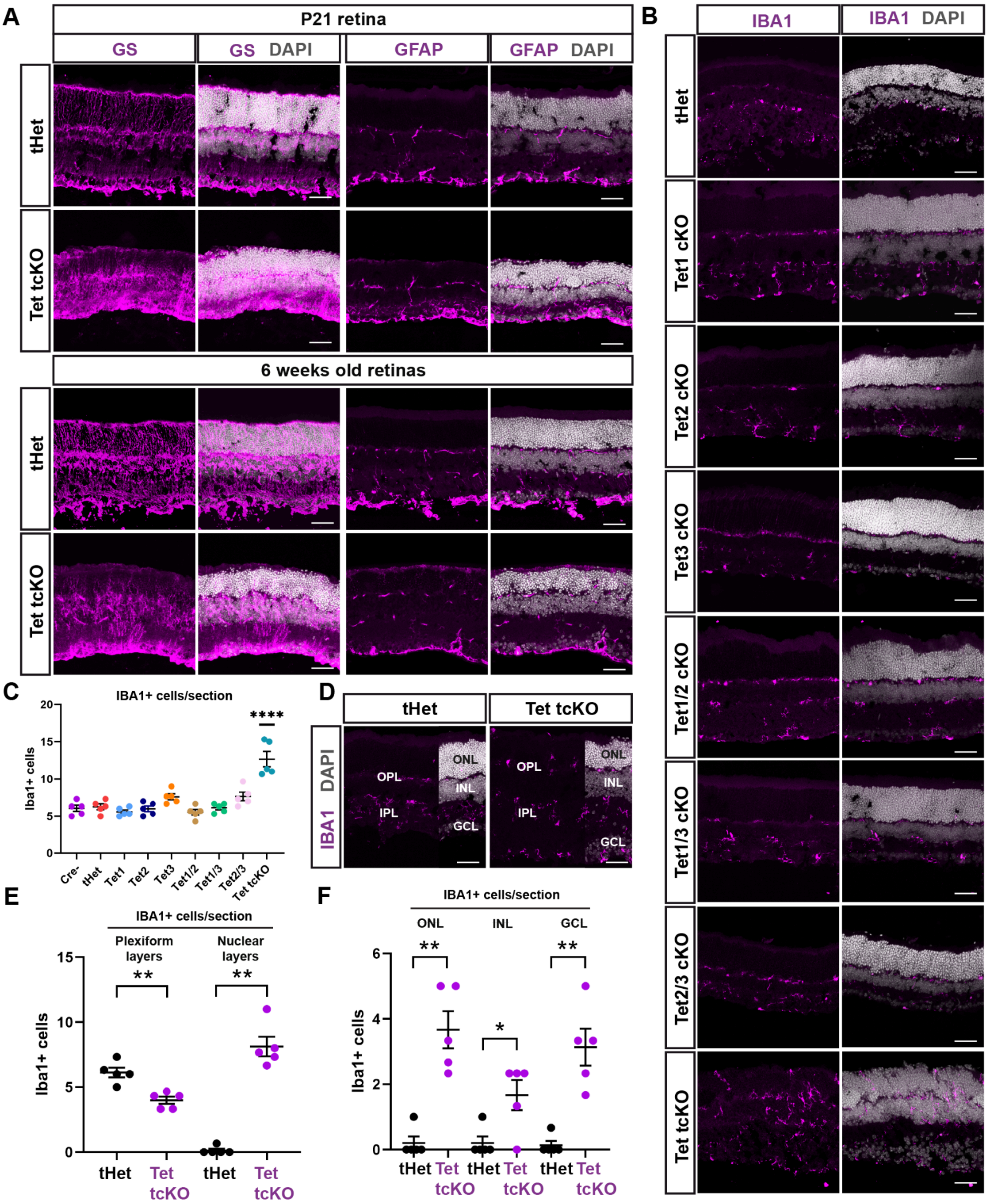


**Supplemental Figure S4. Changes in cell proportions, distribution and morphology of glial cells in TET enzyme mutant retinas. Related to Figure 2.** (A) Immunohistochemistry for Müller glia cells (GS and GFAP) in P21 and 6 weeks old retinas. (B) Immunohistochemistry for microglial cells (IBA1) that show changes in cell proportions of microglial cells when TET enzymes are absent. (C) Results display the mean + SEM for n=5 for each genotype. Statistics are the result of Ordinary One-Way ANOVA, followed by a Dunnett's multiple comparisons test. **** - p<0.0001. (D) Immunohistochemistry for microglial cells (IBA1) that show the comparison between tHet and Tet1/2/3 cKO retinas and the layers that were considered for cell counts in E and F. (E, F) Graphs showing the difference in microglial cell location throughout the different retinal layers comparing tHet and Tet1/2/3 retinas. Results display the mean + SEM for n=5 for each genotype comparing plexiform vs nuclear layers, and the three nuclear layers separately (ONL, INL and GCL). Statistics are the result of an Unpaired t-test or Mann-Whitney test. (E) ** - $p<0.01$. (F) * $p<0.05$; ** - $p<0.01$.

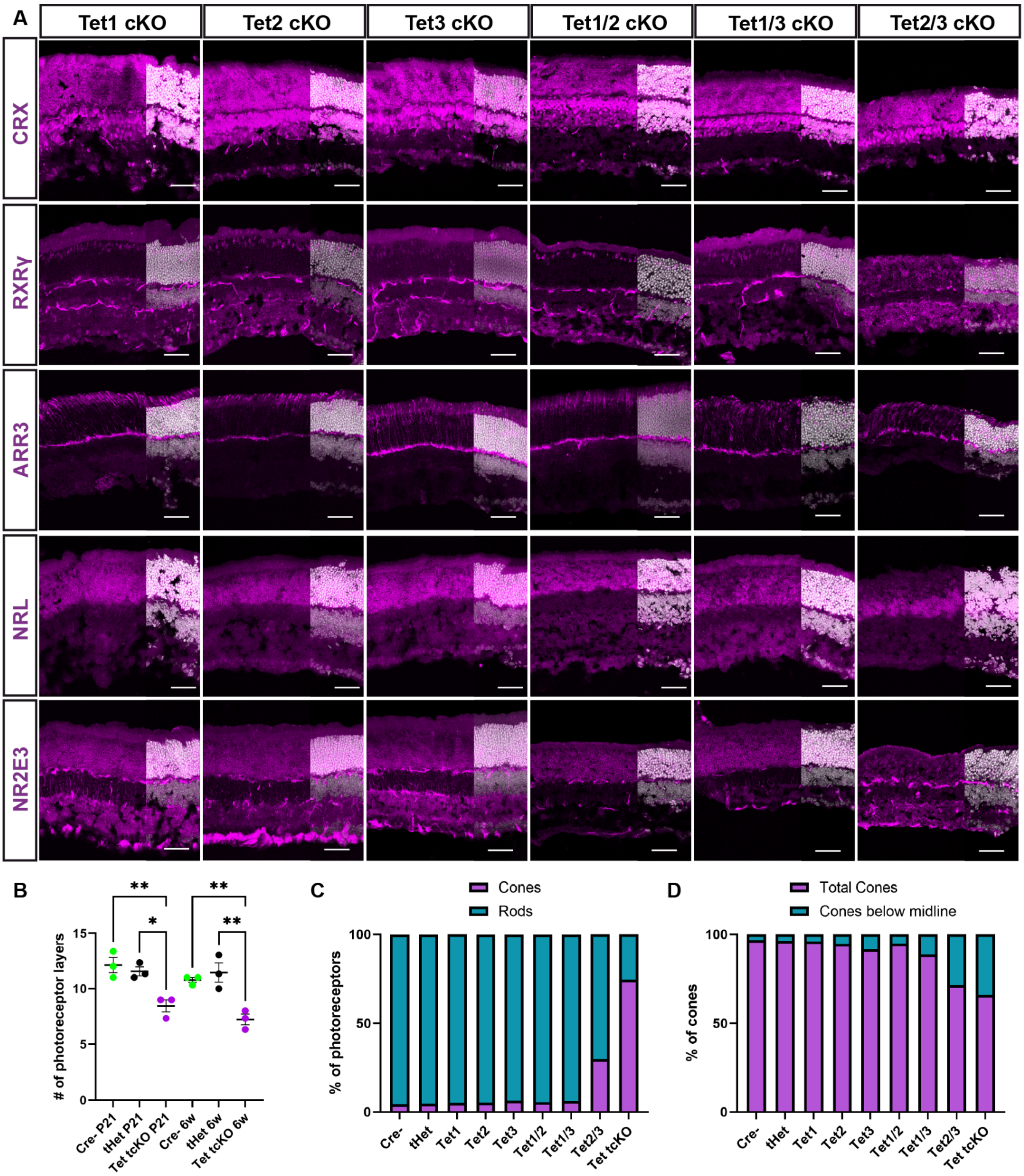


**Supplemental Figure S5. Altered photoreceptor fate specification in TET mutant retinas. Related to Figure 3**. (A) Immunohistochemistry for cone-photoreceptors (CRX, RXRγ, ARR3) and rod-photoreceptors (CRX, NRL and NR2E3) markers that show changes in photoreceptor cells proportions when TET enzymes are absent. (B) Graphs showing the significant decrease in the number of photoreceptor cell nuclei in the ONL comparing Cre-, tHet and Tet tcKO H&E stained retinas both at P21 and 6 weeks. Results display the mean + SEM for n=3 for each genotype. Statistics are the result of One-way ANOVA followed by a Tukey's comparisons test * - p<0.05; ** - p<0.01. (C) Graphs showing the contribution of rod-photoreceptors and cone-photoreceptors to the total number of photoreceptor cells across genotypes. (D) Changes in the location of cone-photoreceptor nuclei in the ONL across genotypes.

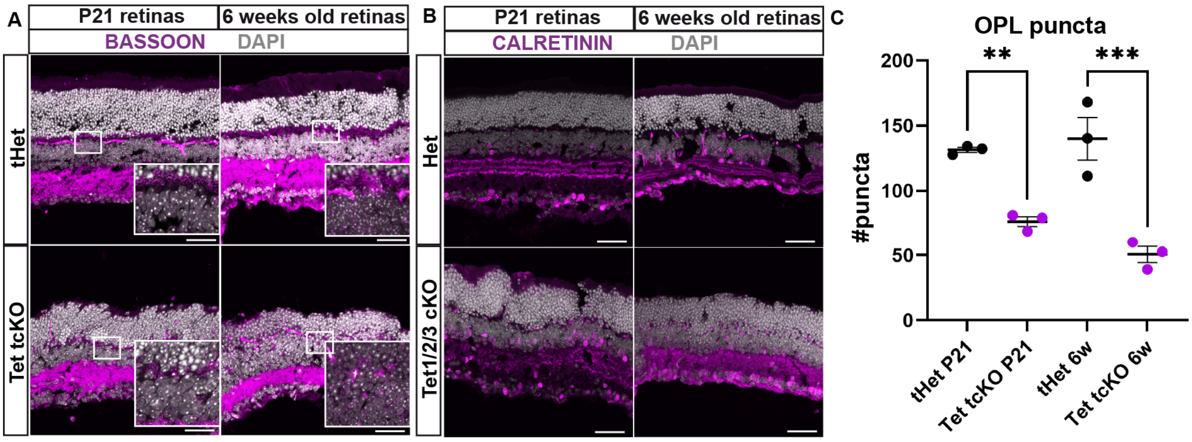


**Supplemental Figure S6**. **Tet tcKO results in altered retinal synapse structures. Related to Figure 1E.** (A and B) Immunohistochemistry for synaptic markers BASSOON and CALRETININ that show the disruption of the OPL and IPL respectively, at P21 and 6 weeks. (C) Graph showing the number of puncta in the OPL labeled by BASSOON. Results display the mean + SEM for n=3 for each genotype. Statistics are the result of Two-Way ANOVA, followed by a Tukey's comparisons test; ** - $p<0.01$; *** - $p<0.001$.

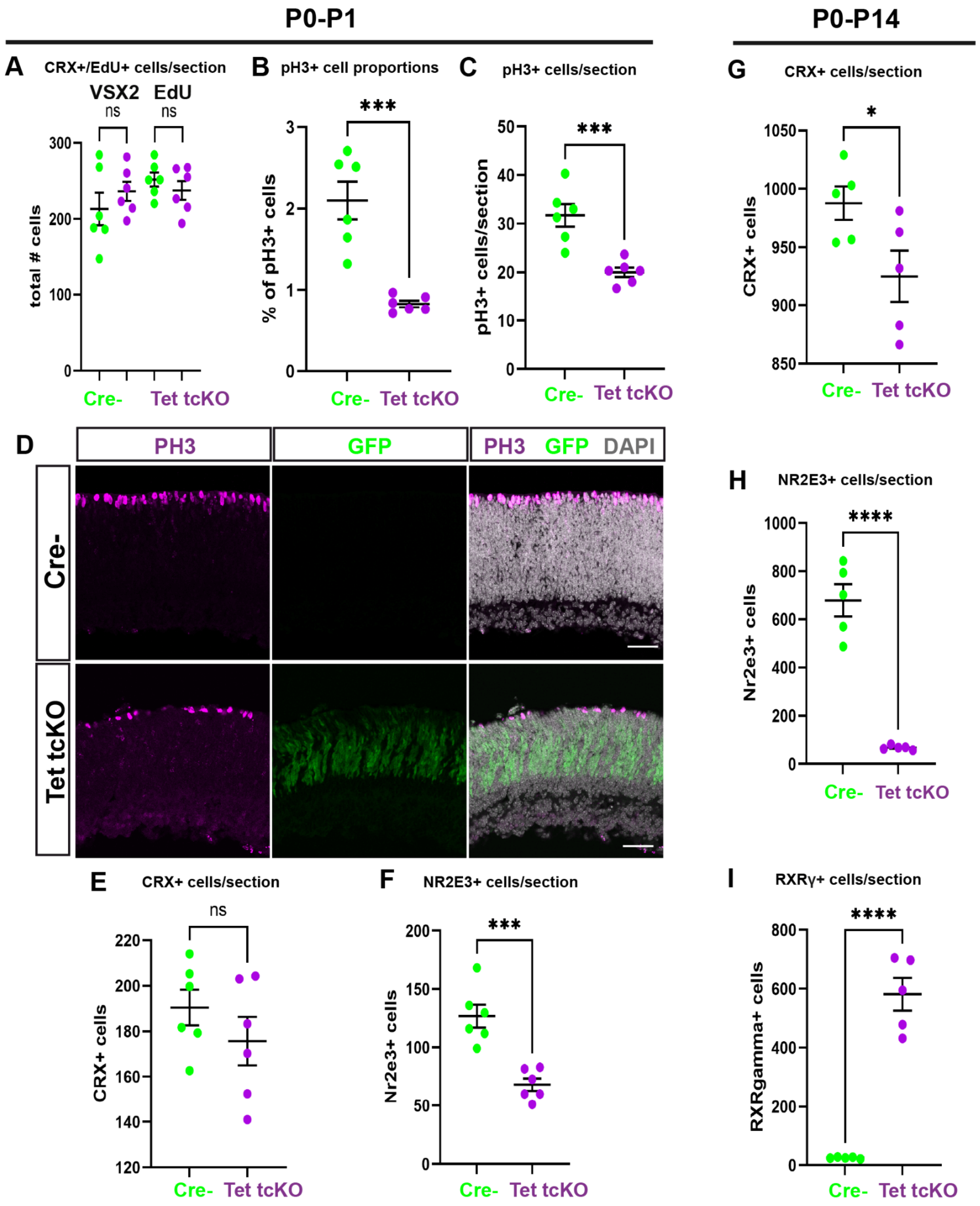


**Supplemental Figure S7. Tet tcKO results in alterations to the total number of progenitor cells and photoreceptors. Related to Figure 4.** (A) Graph showing the total number of VSX2+ and EdU+ cells after P0-P1 EdU pulse. (B and C) Graphs showing the proportion (PH3+/EdU+) and total number of PH3+ cells P0-P1 EdU pulse. (D) Immunohistochemistry showing the labeling of mitotic cells (PH3+ cells). (E and G) Graphs showing the total number of CRX+ cells after P0-P1 and P0-P14 EdU pulses. (F and H) Graphs showing the total number of NR2E3+ and EdU+ cells after P0-P1 and P0-P14 EdU pulses. (I) Graph showing the total number of RXRγ+ cells after P0-P14 EdU pulse. Results display the mean + SEM for n=5 (P0-P14) or n=6 (P0-P1) for each genotype. Statistics are the result of Two-tailed Unpaired t-test; ns: non significant; * - $p<0.05$, *** - $p<0.001$; **** - $p<0.0001$.

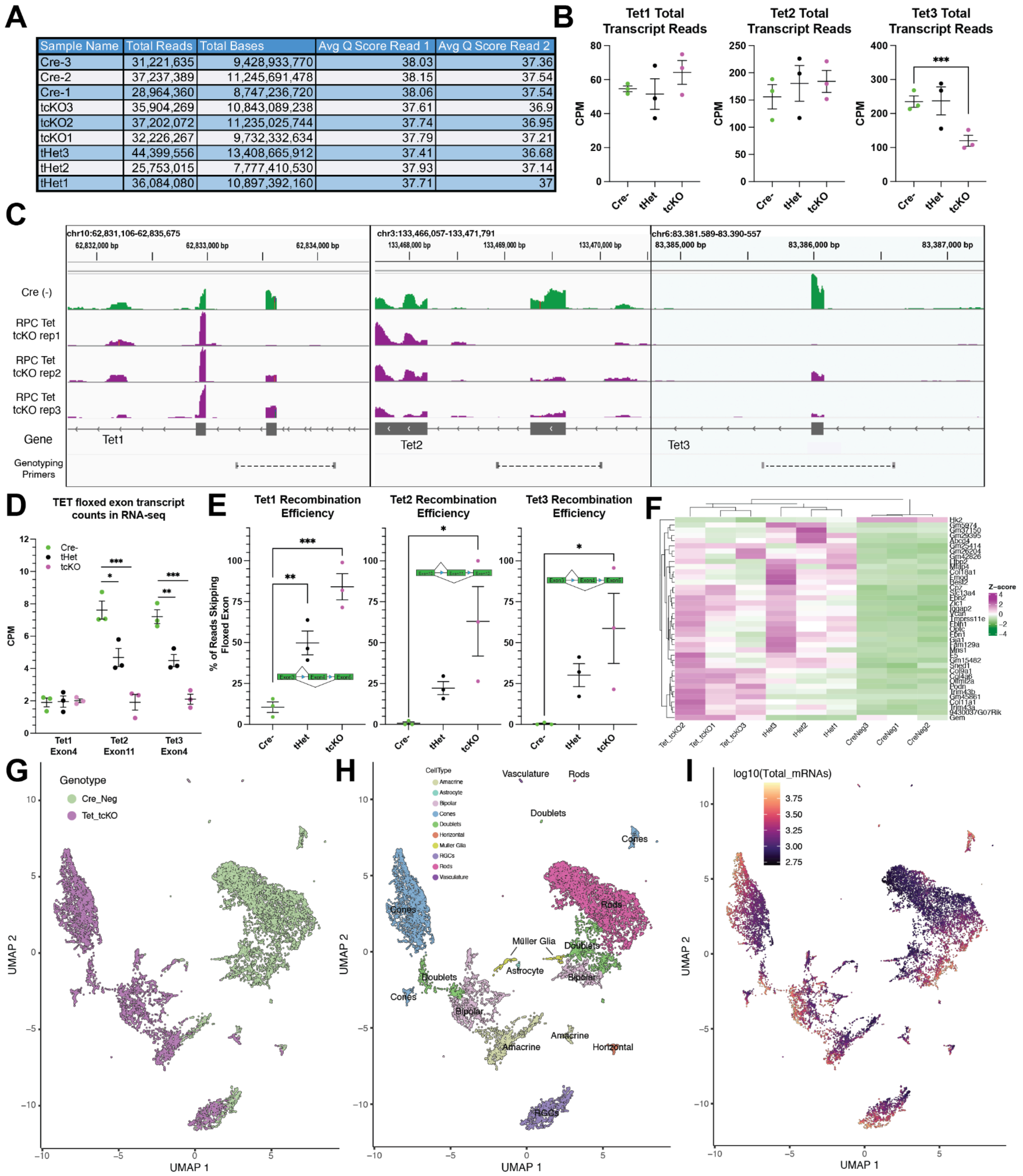

| Sample Name | Total Reads | Total Bases | Avg Q Score Read 1 | Avg Q Score Read 2 |
|---|---|---|---|---|
| Cre-3 | 31,221,635 | 9,428,933,770 | 38.03 | 37.36 |
| Cre-2 | 37,237,389 | 11,245,691,478 | 38.15 | 37.54 |
| Cre-1 | 28,964,360 | 8,747,236,720 | 38.06 | 37.54 |
| tcKO3 | 35,904,269 | 10,843,089,238 | 37.61 | 36.9 |
| tcKO2 | 37,202,072 | 11,235,025,744 | 37.74 | 36.95 |
| tcKO1 | 32,226,267 | 9,732,332,634 | 37.79 | 37.21 |
| tHet3 | 44,399,556 | 13,408,665,912 | 37.41 | 36.68 |
| tHet2 | 25,753,015 | 7,777,410,530 | 37.93 | 37.14 |
| tHet1 | 36,084,080 | 10,897,392,160 | 37.71 | 37 |



**Supplemental Figure S8. Related to Figure 5. Transcriptional profiling of Tet tcKO mutant retinas.** (A) Read depth and quality score metrics for bulk RNAseq samples. (B) Total transcript reads for Tet1 (left), Tet2 (middle) and Tet3 (right) across genotypes. (C) Genome browser tracks indicating read alignment to floxed exons in Cre(-) controls (green) and Tet tcKO replicates (purple) for Tet1 (left), Tet2 (middle) and Tet3 (right). (D) Total counts per million (CPM) for reads aligning to floxed exons across genotypes. (E) Splicing efficiency of floxed exons for Tet1 (left), Tet2 (middle) and Tet3 (right). Splicing efficiency is determined by the ratio of the number of reads that splice into the floxed exon divided by the sum of the splice reads including and excluding the floxed exon. (F) Heatmap of all differentially expressed transcripts (Fold Change > 2; q-value < 0.01) between Cre(-) and tHet RNAseq pairwise comparisons across all RNA-seq replicates. (G-I) UMAP dimension reductions of the full snRNAseq datasets with cells colored by (G) genotype, (H) annotated cell type, and (I) total transcripts detected per cell.

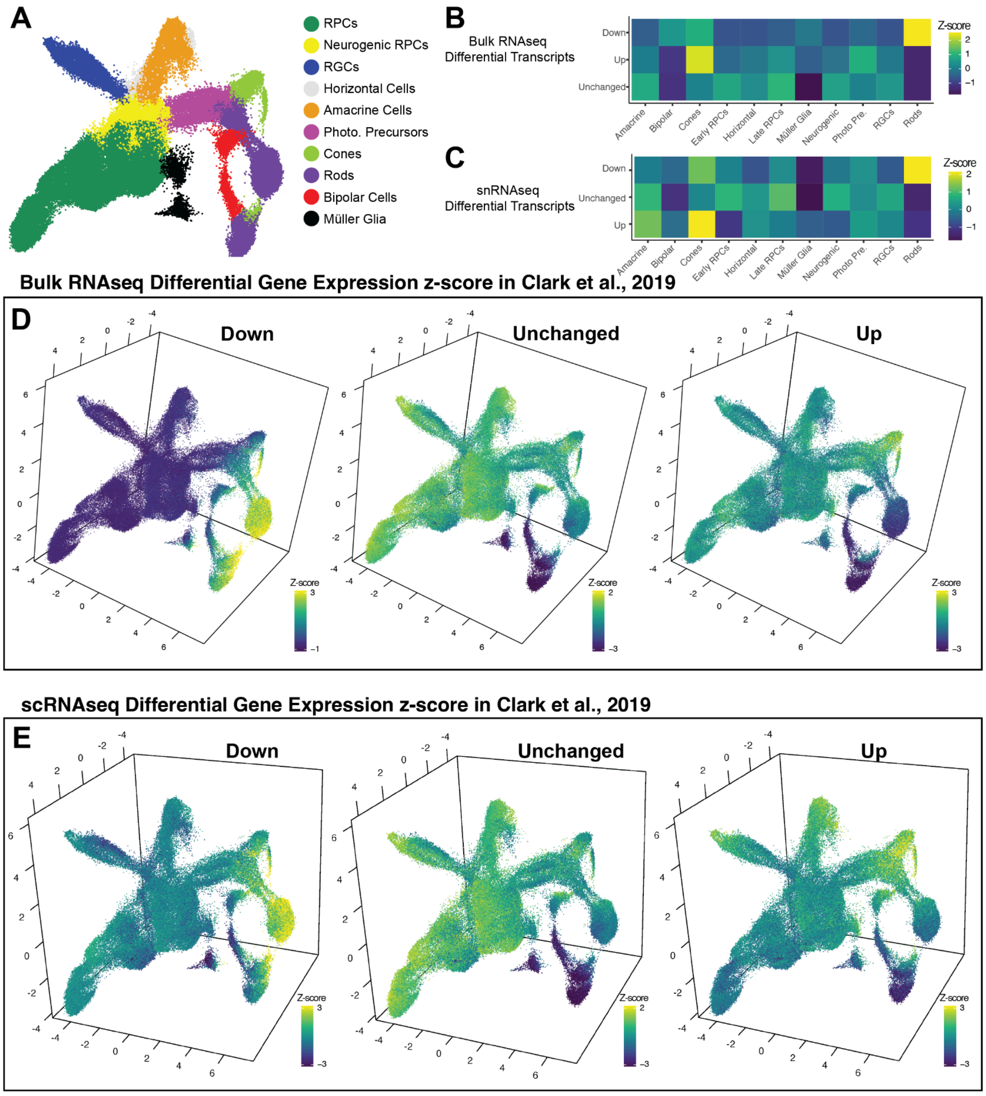


**Supplemental Figure S9 - Related to Figure 5. Gene modules of differentially expressed genes highlight changes in photoreceptor gene expression patterns.** (A) UMAP dimension reduction of the mouse retinal development scRNAseq dataset from Clark, *et al.*, 2019 with cells colored by annotated cell type. (B-C) Heatmaps displaying relative z-scores of differentially expressed transcript gene modules from (B) P21 Tet tcKO RNAseq and (C) P21 Tet tcKO snRNAseq across annotated cell types in the mouse retinal development scRNAseq dataset. (D-E) UMAP dimension reductions of the Clark, *et al.*, 2019 dataset with cells colored by relative z-scores of differentially expressed transcripts as gene modules from (D) P21 Tet tcKO RNAseq and (E) P21 Tet tcKO snRNAseq datasets.

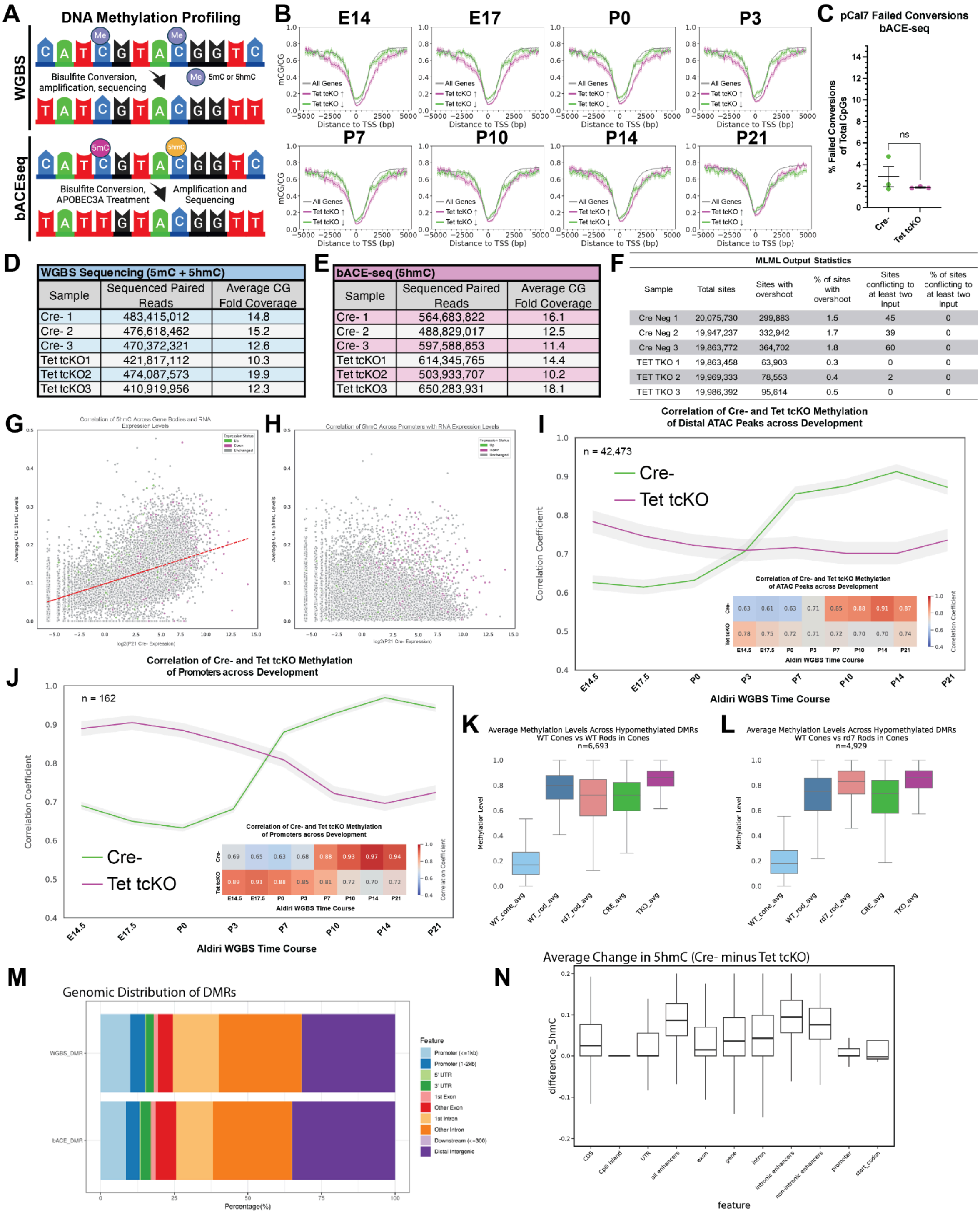


**Supplemental Figure S10. Methylation profiling in control and Tet tcKO retinas. Related to Figure 6.** (A) Graphic displaying results of WGBS and bACE-seq to distinguish 5mC and 5hmC marks. (B) Comparisons of the temporal WGBS methylation patterns across retinal development within the ± 5kb of the TSS for up- and down-regulated transcripts from Tet tcKO RNAseq experiments. (C) Graph highlighting estimated APOBEC3A failed conversion rates utilizing *pCal7* plasmid spiked-in to each bACE-seq reaction. Statistics represent results of a Student's t-test (ns - not significant). (D-E) Summary statistics for WGBS and bACE-seq experiments indicating total reads and average coverage for each sample. (F) MLML Output statistics for distinguishing 5hmC and 5mC marks. (G-H) Scatterplots assessing the correlation RNA expression levels and average 5hmC levels across the (G) gene body or (H) promoter regions for all genes. Individual genes are colored by differential expression in P21 RNAseq comparisons between P21 Cre(-) and Tet tcKO retinas. (I-J) Line plots showing correlation coefficients of P21 Tet tcKO or Cre(-) control WGBS with temporal WGBS methylation profiles across (I) accessible DNA and (J) promoters for ATAC peaks that display a > 10% decrease in methylation levels across development. (K-L) Boxplots of average methylation profiles for differentially hypomethylated regions identified in comparisons between (K) Cones versus Rods and (L) Cones versus *rd7* Rods in P21 Cre(-) and TET tckO retinal samples and sorted cones, rods, and *rd7* rods. (M) Genomic feature distribution for DMRs from WGBS and bACEseq analyses. (N) Boxplots of the change in 5hmC levels (Cre(-) minus Tet tcKO) between P21 Cre(-) and Tet tcKO retinal samples, highlighting the significant loss of enhancer methylation in Tet tcKO retinas.

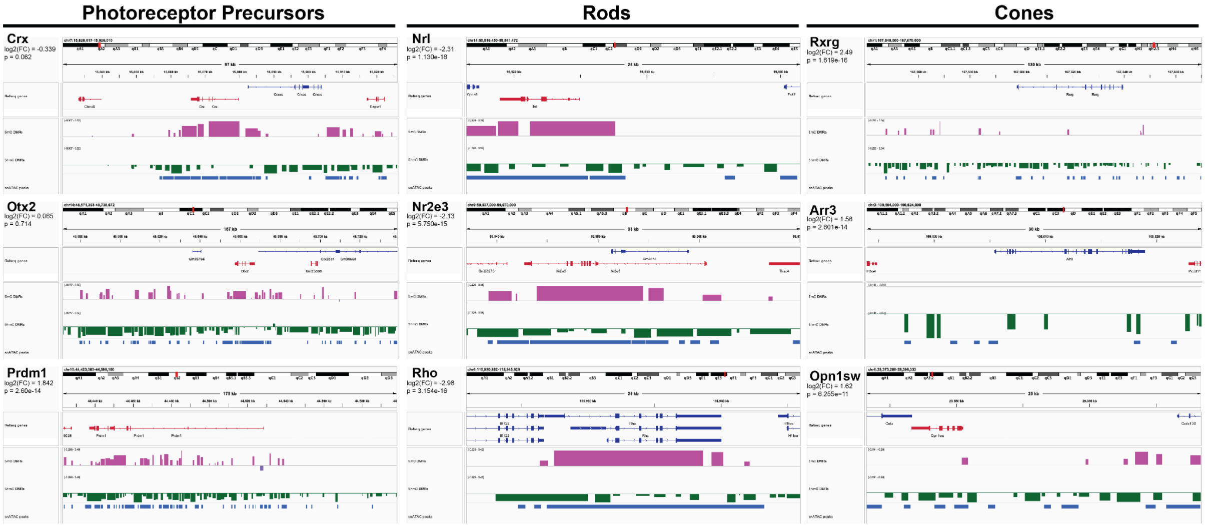


**Supplemental Figure S11 Overlap of 5mC, 5hmC, and snATAC peaks within photoreceptor gene loci. Related to Figure 6.** IGV genome tracks of photoreceptor transcription factor (left), rod photoreceptor (middle) and cone photoreceptor (right) gene loci. DMRs are shown for 5mC (maroon) and 5hmC (green) with the height of the bar representing the average difference in methylation between Tet tcKO and Cre- WGBS and bACE-seq, respectively. 5mC and 5hmC tracks indicate direction of DMR, with bars above or below the grey equivalence lines in each track indicating gain or loss of methylation, respectively. snATAC peak track (bottom, blue) indicates called peaks from the retinal development snATAC-seq studies in Lyu *et al*., 2021.

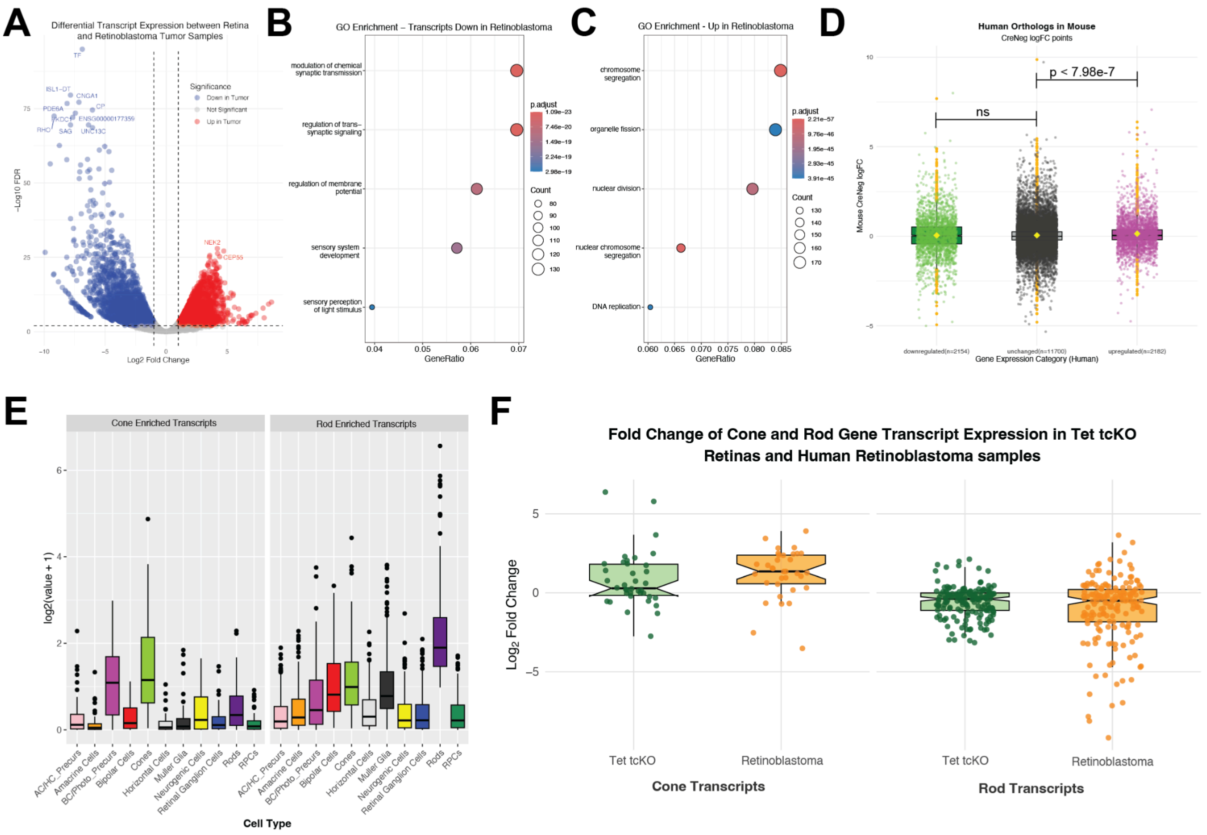

**Supplemental Figure S12 - Related to Figure 7. Retinoblastoma tumors change retinal gene expression patterns.** A) Volcano plot of differentially expressed transcripts between normal retina and retinoblastoma tumor samples with gene names listed for most significant transcript expression differences. (B and C) GO pathway analysis for significant (B) down-regulated and (C) up-regulated transcripts from RNAseq between normal retina and retinoblastoma tumor samples. (D) Relative fold change of mouse orthologs in Tet tcKO versus Cre- retinal samples for retinoblastoma differentially expressed transcripts. Statistics represent the results of pairwise Wilcoxon Rank Sum analyses in comparison to unchanged control transcripts. (E) Average expression of cone-enriched (left) and rod-enriched (right) transcripts from the human single-cell RNAsequencing experiments performed in Lu *et al.*, 2020. (F) Relative fold change of cone (left) and rod-enriched (right) transcripts between mouse P21 Tet tcKO and Cre- (green) or human retinoblastoma tumor and normal retina (orange) RNAseq samples.

**SUPPLEMENTAL TABLES:**

**Table S1.** List of Antibodies used in these studies, including concentrations at which antibodies were used in immunohistochemistry experiments

**Table S2.** RNA-seq results from P21 Cre-, tHet, and Tet tcKO retinas. Table includes results of differential expression analysis

**Table S3.** GO Pathway analysis results for differentially expressed transcripts between P21 Cre- and Tet tcKO retina RNA-seq analysis.

**Table S4.** Results of differential expression analysis across genotypes in snRNA-seq of P21 Cre- and Tet tcKO retinal samples.

**Table S5.** Differential methylation (5mC + 5hmC) analysis of WGBS on P21 Cre- and Tet tcKO whole retinas. Results indicate the genomic location of DMRs, the number of CpGs present within the genomic locus, and the mean methylation levels across Cre- and Tet tcKO samples.

**Table S6.** Differential methylation analysis (5hmC) of bACE-seq experiments on P21 Cre- and Tet tcKO whole retinas. Results indicate the genomic location of 5hmC DMRs, the number of CpGs present within the genomic locus, and the mean 5hmC levels across Cre- and Tet tcKO samples.

**Table S7.** RNA-seq results of human retinoblastoma tumor samples and normal retina controls. Table includes results of differential expression analysis.

**Table S8.** Transcripts that show enrichment in either rods and cones from the human developmental single-cell RNA-sequencing dataset from Lu *et al*., 2020.
